# TorsinA is essential for the timing and localization of neuronal nuclear pore complex biogenesis

**DOI:** 10.1101/2023.04.26.538491

**Authors:** Sumin Kim, Sébastien Phan, Thomas R. Shaw, Mark H. Ellisman, Sarah L. Veatch, Sami J. Barmada, Samuel S. Pappas, William T. Dauer

## Abstract

Nuclear pore complexes (NPCs) regulate information transfer between the nucleus and cytoplasm. NPC defects are linked to several neurological diseases, but the processes governing NPC biogenesis and spatial organization are poorly understood. Here, we identify a temporal window of strongly upregulated NPC biogenesis during neuronal maturation. We demonstrate that the AAA+ protein torsinA, whose loss of function causes the neurodevelopmental movement disorder DYT-TOR1A (DYT1) dystonia, coordinates NPC spatial organization during this period without impacting total NPC density. Using a new mouse line in which endogenous Nup107 is Halo-Tagged, we find that torsinA is essential for correct localization of NPC formation. In the absence of torsinA, the inner nuclear membrane buds excessively at sites of mislocalized, nascent NPCs, and NPC assembly completion is delayed. Our work implies that NPC spatial organization and number are independently regulated and suggests that torsinA is critical for the normal localization and assembly kinetics of NPCs.

## INTRODUCTION

Nuclear pore complexes (NPCs) are large, evolutionarily conserved structures that serve as the gateway between the nucleus and cytoplasm, allowing passive diffusion of small molecules and facilitating nucleocytosolic transport of proteins and RNA^1,2^. Regulation of NPC biogenesis and function is critical for coordinating information transfer between the nucleus and cytoplasm, allowing cells to dynamically respond to internal and external cues. Neurons are heavily dependent on NPC function; neuronal plasticity requires transport of signaling molecules and transcription factors^3–6^ and mRNA export for local translation is essential for several neurodevelopmental processes^7–11^. Accordingly, NPC defects are present in several nervous system diseases^12–16^ and mutations in nucleoporins cause early-onset neurological illness^17,18^. Indeed, neurons face a considerable challenge in maintaining proper nuclear function throughout their lifetime. Whereas mitotic cells disassemble and recreate the nucleus with each division, neurons must manage nuclear pore number and localization within a closed interphase nucleus. Neuronal NPCs exhibit minimal turnover and nucleoporins (Nups) that constitute the NPC are among the longest-lived proteins^19,20^, underscoring the unique challenge neurons face in regulating NPC formation and function. Despite the biological and clinical importance of these events, little is understood about neuronal NPC biogenesis. Furthermore, mechanisms underlying NPC number, organization, and dynamics in both neuronal and non-neuronal systems remain elusive.

TorsinA appears to lie at the intersection of neuronal NPC biogenesis and neurodevelopmental disease. TorsinA is a AAA+ protein that resides in the endoplasmic reticulum (ER)/nuclear envelope (NE) lumen^21–29^. The neurodevelopmental movement disorder DYT-TOR1A (DYT1) dystonia is caused by an in-frame 3-bp deletion in the TOR1A gene that encodes a ΔE-torsinA mutant protein^30,31^. Several observations demonstrate that the NE is an active site of torsinA activity. A “substrate trap” mutant of torsinA primarily localizes to the NE^24,25^ and perturbing torsinA levels causes changes in the LINC (linker of nucleoskeleton and cytoskeleton) complex^32,33^. Mislocalization of nuclear membrane proteins is also observed in *C. elegans* germ cells lacking the torsin homolog OOC-5, which additionally exhibit asymmetric plaques of Nups^34^.

TorsinA-knockout (KO) or homozygous ΔE mutant mice develop abnormal NE evaginations or “blebs” exclusively in post-migratory maturing neurons^35^, establishing ΔE as a loss-of-function (LOF) mutation. Biochemical studies are consistent with a LOF effect of the ΔE mutation^36,37^. NE blebs are inner nuclear membrane (INM) outpouchings that project into the NE lumen. These blebs occur transiently, emerging during early postnatal CNS maturation and subsequently resolving without causing cell death^35,38,39^. Non-neural tissue and mitotic cell lines lacking torsinA do not exhibit NE abnormalities^40–42^, highlighting the unique requirement for torsinA function in neurons. Indeed, in vivo studies demonstrate that torsinA function is uniquely essential during early postnatal neural development^43^, emphasizing the importance of probing torsinA function in this context.

The INM budding characteristic of NE blebs suggests a potential link to interphase NPC assembly. During this process, new NPCs are inserted *de novo* into an intact nuclear membrane via INM budding and subsequent fusion with the outer nuclear membrane (ONM)^44–46^. Dissimilar to the budding observed in intermediate stages of normal NPC assembly, NE blebs do not fuse with the ONM and continue to enlarge into the NE lumen. Maturing torsinA-KO neurons also develop mislocalized clusters of nucleoporins which, unlike NE blebs, persist into adulthood in DYT1 mouse models^13,43^. These clusters contain Nup153, an NPC component incorporated during initial steps of biogenesis, but lack the later-recruited cytoplasmic nucleoporin Nup358 (added to the NPC after INM-ONM fusion^44^). The lack of INM-ONM fusion in NE blebs and the absence of Nup358 in nucleoporin clusters suggests that these two phenotypes may be related and implicates a potential role for torsinA in NPC biogenesis. Indeed, mitotic cells lacking multiple torsin protein family members (torsinA, torsinB, torsin2A, torsin3A) develop NE blebs that co-localize with NPC components and are downregulated by blocking interphase NPC assembly^40,47^. Nucleoporin distribution appears normal in these cells^42,47^, however, precluding their use for exploring this striking, disease-relevant phenotype. Little is known about these clusters, including their relationship to NE blebs, whether they contain formed (but mislocalized) NPCs or aggregated nucleoporins, and how they relate to NPC assembly.

Here, we identify a novel role for torsinA in the localization and timing of neuronal NPC biogenesis. We present a quantitative analysis of neuronal NPC biogenesis and find that NPC formation is strongly upregulated during a distinct neurodevelopmental window. To study physiological NPC biogenesis in neurons, we created a novel line of knock-in mice harboring HaloTag-Nup107 allele. Pulse-chase studies in primary neurons derived from these mice demonstrate that torsinA is essential for the normal localization of nascent NPCs. Abnormal clusters containing mislocalized NPCs arise simultaneously with the emergence of NE blebs in torsinA-KO neurons. Ultrastructural evidence from electron tomography demonstrates that NE blebs form at the sites of NPC biogenesis and represent a stalled intermediate step of NPC assembly. While abnormal NPC clusters persist, NE blebs resolve in mature neurons. Completion of new NPCs is delayed, but not permanently halted, in torsinA-KO neurons, indicating that torsinA is not essential for INM-ONM fusion as previously believed. Our work advances understanding of NPC spatial organization and newly highlights a critical role for torsinA in NPC biogenesis during a discrete developmental window.

## RESULTS

### NPC biogenesis is upregulated during neuronal maturation

To explore neuronal NPC biogenesis, we used structured illumination microscopy (SIM) to characterize NPC density in cortical primary neurons. Primary neurons from P0 pups were cultured and fixed at days in vitro (DIV) 4, 6, 8, 10, 14, 18, and 24 and labeled with Nup153 and Nup98 antibodies. In immature neurons^48^ (i.e., DIV4 and DIV6), both nucleoporins (Nups) appeared sparse. As the neurons matured, the density of Nup153 and Nup98 puncta increased, culminating in a dense yet even distribution of NPCs across the nuclear envelope (DIV18 and DIV24; Figure 1A, Figure S1A). We developed an image analysis pipeline to identify and localize the peaks of Nup puncta to quantitatively measure Nup density and distances (Figure S1B). Peaks from Nup153 and 98 puncta appeared to mostly overlap, suggesting that the antibodies used in this approach labeled NPCs rather than non-NPC pools of nucleoporins. The density of Nup153 and Nup98 puncta gradually increased over two weeks of neuronal maturation from approximately 7 puncta/μm^2^ at DIV4 to 14 puncta/μm^2^ at DIV24 (Figure 1B, 1C). Nuclear area did not change significantly during this time (Figure S1C), indicating that the increased Nup density is due to *de novo* formation of new NPCs. We used coordinates of identified peaks to calculate the nearest neighbor distance for each punctum (Figure S1D, S1E). As expected, few puncta at any timepoint measured have a nearest neighbor within 120nm, the diameter of a single NPC^49,50^. The percentage of Nup153 and Nup98 puncta with a nearest neighbor within a two-pore diameter distance (240nm) increased with time, while the percentage of puncta with a nearest neighbor far away decreased. Consistent with increasing NPC density, the proportion of Nup puncta with multiple proximal neighbors increased as neurons matured (Figure 1D, 1E). Together, these data demonstrate that NPC biogenesis is upregulated during neuronal maturation.

**Figure 1:**
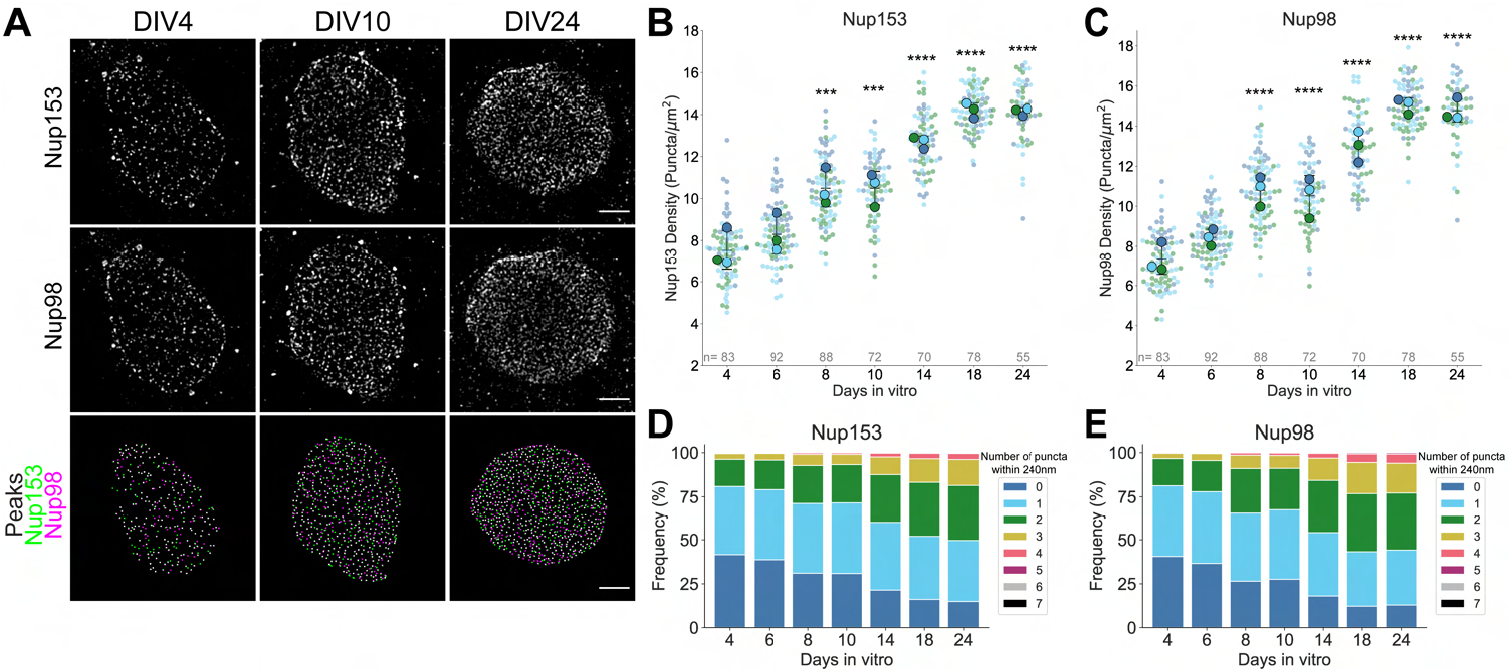
Nuclear pore complex biogenesis is upregulated during neuronal maturation. **A**. Structured illumination microscopy (SIM) images of primary neurons aged DIV4, 10, and 24 labeled with anti-Nup153 and anti-Nup98 antibodies. Bottom row shows identified Nup153 and 98 peaks. Scale bar = 2μm. **B**. Superplots of Nup153 density (puncta/μm^2^) over time in primary neurons. Plots show mean ± SD, with color coding indicative of biological replicates. Repeated measures one-way ANOVA with Dunnett’s multiple comparisons test was performed, with DIV4 as the reference condition. ***P=0.0006, ****P<0.0001. Total number of analyzed nuclei are represented as “n”. **C**. Superplots of Nup98 density (puncta/μm^2^) over time in primary neurons. Repeated measures one-way ANOVA with Dunnett’s multiple comparisons test was performed, with DIV4 as the reference condition. ****P<0.0001. **D, E**. Frequency distribution of Nup153 (D) and Nup98 (E) puncta found within two-pore diameter (240nm) distance. For each identified Nup153 and Nup98 peak, the number of neighboring puncta within a radius of 240nm was calculated. Results from three biological replicates were combined at each timepoint.

The acquisition of SIM images and subsequent processing time are not well suited for large-scale analysis of cultured neurons. We therefore pursued the alternative strategy of measuring nuclear rim intensity of NPC components with confocal microscopy to further explore NPC biogenesis. Consistent with our SIM data, there was an approximately twofold increase in nuclear rim intensity of Nup153 and Nup98 between DIV4 and DIV10 (Figure S2A-S2C). Additionally, we used antibodies against FG-Nups (Nups containing Phe-Gly motifs; mab414) and Nup210 (a transmembrane nucleoporin) to assess whether other Nups were similarly upregulated, as would be expected with increased NPC biogenesis. Between DIV4 and DIV10, the nuclear rim intensity of mab414 and Nup210 increased twofold (Figure S2D-S2F), demonstrating upregulated density of multiple NPC components, including those spanning the nuclear basket, inner ring, and transmembrane subunits.

### TorsinA is essential for uniform NPC distribution but not upregulation of NPC density

We reported previously that torsinA deletion results in long-lasting abnormal clusters of nucleoporins in neurons^13^. We asked whether neuronal NPC biogenesis is affected in the absence of torsinA and whether formation of abnormal nucleoporin clusters coincides with increased NPC biogenesis. We fixed maturing WT and torsinA-KO neurons at DIV4, 6, 8, 10, and 14 and labeled NPCs using anti-Nup153 and anti-Nup98 antibodies. At DIV4, Nup153 and Nup98 were sparsely localized in both WT and torsinA-KO neurons such that their distribution was indistinguishable between genotypes (Figure 2A, 2B). While nucleoporin organization remained uniform in maturing WT neurons, DIV6 torsinA-KO neurons exhibited small clusters that multiplied and enlarged over time (Figure 2A, 2B). Mature torsinA-KO neurons displayed strikingly large clusters, contrary to the dense but uniform distribution observed in mature WT neurons. We conducted an autocorrelation analysis to quantitatively characterize changes in nucleoporin spatial correlation across developmental age (Figure S3A, S3B). Autocorrelation analysis conveys two types of information: the density of particles is roughly inversely correlated to the amplitude of the autocorrelation curve, and autocorrelation values indicate dispersion (<1) or clustering (>1). The decreasing amplitude of the autocorrelation curve reflected increasing density over time for both WT and torsinA-KO neurons, while a broadening of the curve for torsinA-KO neurons indicated growing clusters (Figure S3A). We normalized autocorrelation to total density at each timepoint to account for amplitude dependence on density. In WT neurons, normalized autocorrelation curves at all timepoints exhibited a dip <1 around 200nm indicating dispersion of nucleoporins at this distance (Figure S3B). In contrast, the normalized autocorrelation increased over time for torsinA-KO neurons, reflecting progressively worsening clusters.

**Figure 2:**
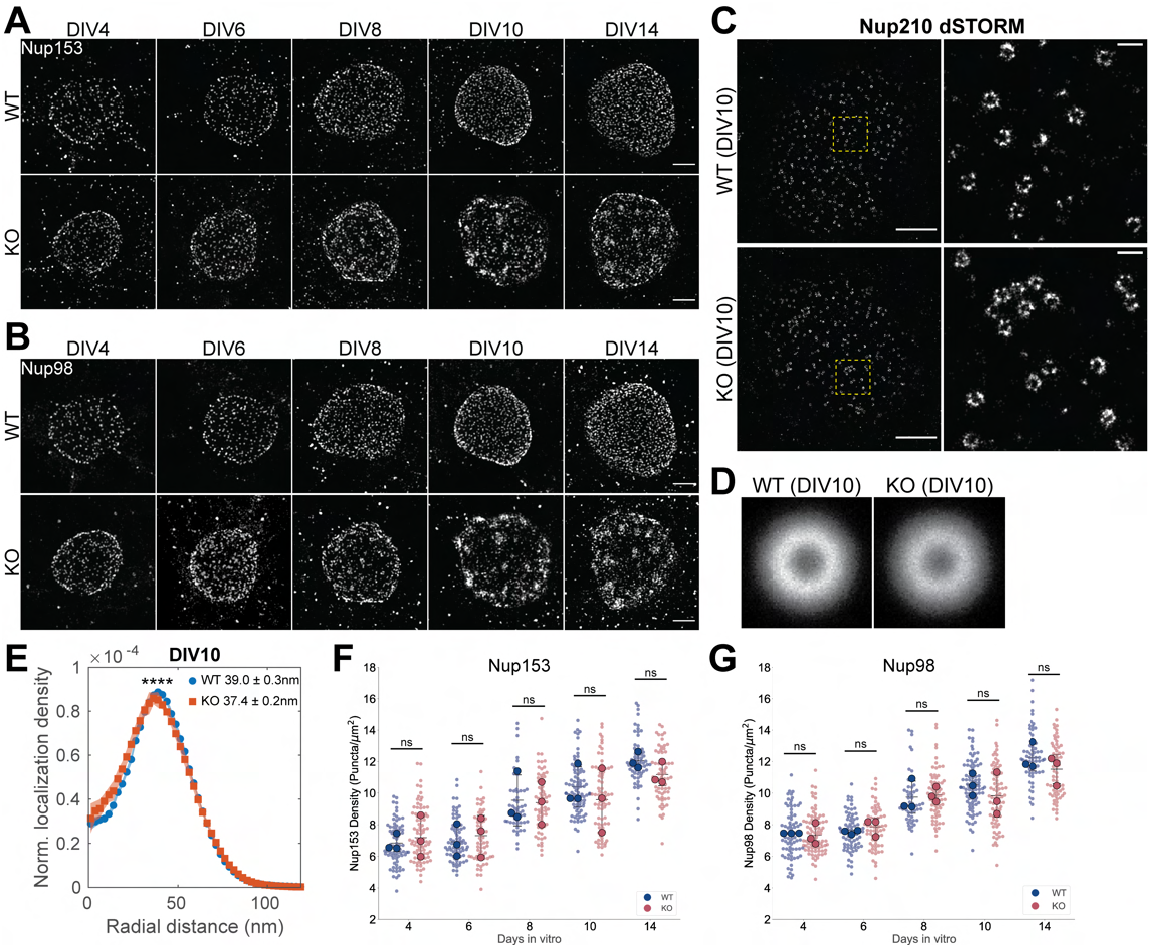
TorsinA is essential for uniform NPC distribution but not upregulation of NPC biogenesis. **A, B**. SIM images of WT and torsinA-KO primary neurons aged DIV4, 6, 8, 10, and 14 labeled with anti-Nup153 (A) and anti-Nup98 (B) antibodies. Scale bar = 2μm. **C**. dSTORM images of Nup210-labeled NPCs in DIV10 WT and torsinA-KO neurons. Scale bar = 2nm. Right panels show zoomed in view of boxed regions. Scale bar of right panels = 200nm. **D**. Averaged aligned DIV10 WT and torsinA-KO pores. **E**. Normalized localization density along radial distance in averaged WT and KO pores. Plots show mean ± SEM from eight bootstrapping rounds with 250 randomly selected pores each. ****P<0.0001. **F, G**. Nup153 (F) and Nup98 (G) density (puncta/μm^2^) in maturing primary neurons. Plots show mean ± SD. Timepoints from each replicate were matched and repeated measures two-way ANOVA with Sidak’s multiple comparisons test was performed to compare means between genotypes. ns, not significant.

While our SIM data demonstrate that nucleoporins form clusters at the nuclear envelope as torsinA-KO neurons mature, they do not address whether nucleoporins are recruited to NPCs or form unstructured aggregates. We therefore asked whether abnormal nucleoporin clusters contain structures resembling normal NPCs using direct stochastic optical reconstruction microscopy (dSTORM). We labeled DIV10 neurons using a Nup210 antibody because Nup210 subunits are spaced farther apart than those of Nup98 and Nup153, enabling easier detection of individual subunits. WT neurons showed clear, uniformly distributed NPC structures containing central channels (Figure 2C). Similar NPC structures containing central channels were observed in torsinA-KO DIV10 neurons but were localized in clusters. We again used autocorrelation to analyze spatial correlation between NPCs. In WT neurons, the autocorrelation value was <1 between 150nm and 400nm, indicating dispersion (Figure S3C). In contrast, autocorrelation >1 was observed in torsinA-KO neurons over the same distance range, reflecting clustering. To further analyze spatial organization and structure of individual NPCs, we segmented individual NPCs (see Methods). Plotting the nearest neighbor distance between centroids of segmented NPCs in WT neurons yielded two distinct peaks, indicating regular spatial organization of NPCs (Figure S3D). In torsinA-KO neurons, however, the second peak was absent, reflecting abnormal NPC spacing. We then asked whether WT and torsinA-KO NPCs exhibit structural differences. We aligned and averaged segmented NPCs (Figure 2D, see Methods) and calculated their radii (Figure 2E). The average WT NPC exhibited a radius of 39nm +/- 0.3nm, while the average KO NPC had a smaller radius of 37.4 +/- 0.2nm. Despite this slight difference in NPC channel, these data clearly indicate that abnormal nucleoporin clusters in torsinA-KO neurons represent aberrantly localized clusters of NPCs rather than unstructured aggregates. Herein we refer to nucleoporin clusters as NPC clusters.

Based on our characterization of the identity of NPC clusters, we calculated NPC density in WT and torsinA-KO neurons from SIM images (Figure 2A, 2B), incorporating puncta identified from clusters. Despite striking differences in spatial organization, there was no significant difference in average NPC density between WT and torsinA-KO neurons at any age (Figure 2F, 2G, Figure S3E, S3F), indicating that maturing WT and torsinA-KO neurons similarly upregulate NPC biogenesis. Nuclear area and overall shape also did not differ significantly between WT and torsinA-KO neurons (Figure S3G, S3H). Consistent with these findings, expansion microscopy to label Nup210 and LaminA/C showed no deformities in the overall nuclear shape of torsinA-KO neurons (Figure S3I). Furthermore, abnormal NPC clusters did not correspond to any local folds or dips in the lamin meshwork. Collectively, these data strongly suggest that torsinA is essential for the proper spatial organization, but not total density, of NPCs.

### Novel HaloTag-Nup107 knock-in allele enables labeling of endogenous NPC dynamics

To dissect mechanistically how NPC clusters form, we developed a novel tool to track endogenous NPC biogenesis. We generated a knock-in mouse line in which HaloTag^51^ is fused to the N-terminus of endogenous Nup107 (Figure 3A). Nup107 was chosen as prior literature demonstrates that N-terminally tagged 3x EGFP-Nup107 maintains normal localization and dynamics^52,53^. We validated this HaloTag-Nup107 (Halo-Nup107) knock-in mouse line using the JF646 HaloTag ligand to label endogenous Nup107 protein in DIV10 cortical primary neurons (Figure 3B). We observed a gene dose-dependent increase in fluorescence intensity at the nuclear membrane in neurons derived from *Nup107*^*+/+*^, *Nup107*^*KI/+*^, *Nup107*^*KI/KI*^ mice, indicating that the fusion protein is incorporated into nuclear pores. We further assessed the knock-in by probing the levels of the Nup107-HaloTag fusion protein in P0 cortical lysates. We observed a single band corresponding to Nup107 in *Nup107*^*+/+*^ lysate, two bands representing both Nup107 and HaloTag-Nup107 in *Nup107*^*KI/+*^ lysate, and a single band indicative of HaloTag-Nup107 in *Nup107*^*KI/KI*^ lysate (Figure S4A). *Nup107*^*KI/KI*^ mice were indistinguishable from their WT littermate controls and we observed the expected Mendelian ratios of genotypes from *Nup107*^*KI/+*^ intercross (Figure S4B), indicating that the HaloTag did not exert any toxicity.

**Figure 3:**
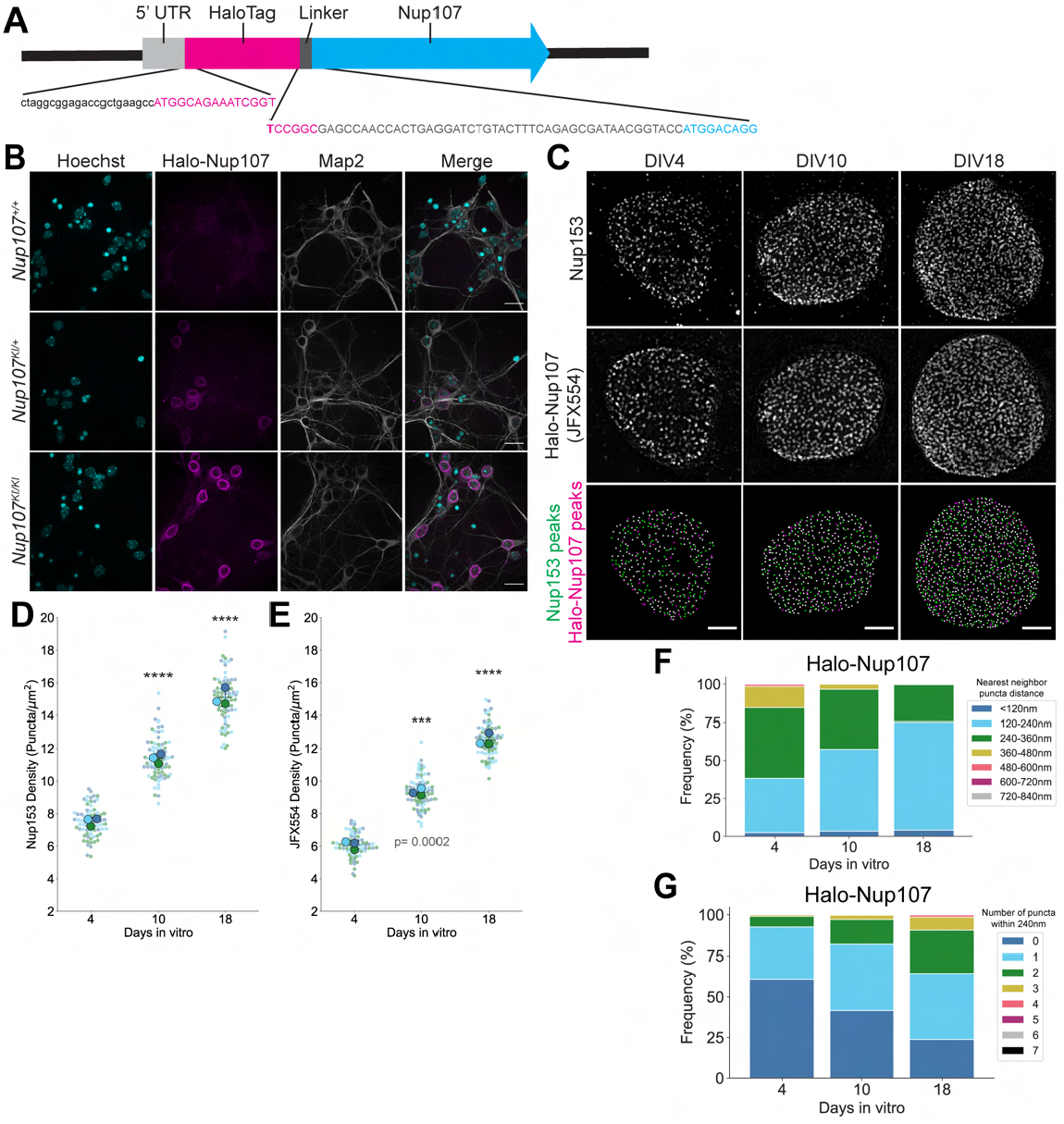
Neurons bearing a novel HaloTag-Nup107 knock-in allele demonstrate endogenous NPC upregulation. **A**. Schematic of HaloTag-Nup107 fusion, including 5’ untranslated region (UTR, light grey), HaloTag open reading frame (magenta), flexible linker (dark grey) and Nup107 coding sequence (cyan). **B**. Confocal images of DIV10 neurons derived from *Nup107*^*+/+*^, *Nup107*^*KI/+*^, and *Nup107*^*KI/KI*^ mice labeled with JF646 HaloTag ligand. Scale bar = 20μm. **C**. SIM images of *Nup107*^*KI/KI*^ primary neurons at DIV4, 10, and 18 labeled with JFX554 HaloTag ligand and anti-Nup153 antibody. **D, E**. Superplots of Nup153 (D) and JFX554 (E) puncta density in primary neurons aged DIV4, 10, 18. Plots show mean ± SD, with color coding indicative of biological replicates. Repeated measures one-way ANOVA with Dunnett’s multiple comparisons test was performed, with DIV4 as the reference condition; ****P<0.0001. Number of analyzed nuclei are represented as “n”. **F**. Frequency distribution of nearest neighbor distances of JFX554 puncta. For each JFX554 peak, the distance to its nearest neighbor was calculated. Results from three biological replicates were combined. **G**. Frequency distribution of JFX554 puncta within two-pore diameter (240nm) distance. For each JFX554 peak, the number of neighboring puncta within a radius of 240nm was calculated. Results from three biological replicates were combined.

We examined the localization and density of HaloTag-Nup107 in developing neurons by labeling DIV4, 10, and 18 *Nup107*^*KI/KI*^ neurons with the JFX554 HaloTag ligand and staining them with Nup153 antibody. SIM imaging of single NPCs demonstrated an approximately twofold increase in Nup density from DIV4 to DIV18 despite similar nuclear envelope area (Figure 3C-3E, Figure S4C). Overlay of Nup153 peaks with JFX554 puncta and vice versa showed a high level of colocalization (Figure S4D, S4E). Nearest neighbor distances of HaloTag-Nup107 puncta decreased throughout neuronal maturation while the number of neighboring puncta within 240nm increased, reflecting the increase in density and decrease in pore-pore distances as neurons mature (Figure 3F, 3G). These data validate the HaloTag-Nup107 mouse model as a novel tool for studying endogenous NPC dynamics and confirm that maturing neurons exhibit strongly upregulated NPC biogenesis.

### Sites of NPC biogenesis are abnormal in torsinA-KO neurons

To elucidate how NPC clusters arise in the absence of torsinA, we performed a pulse-chase experiment using HaloTag-Nup107 neurons to label NPCs formed at different stages of neuronal maturation. We crossed *Nup107*^KI/KI^ mice with the *Tor1a*^+/-^ mouse line^35^ and did not observe any deviations from Mendelian genotypic ratios in *Tor1a*^+/-^ ;*Nup107*^*KI/KI*^ intercross (Figure S5A). Neurons from WT (*Tor1a*^+/+^;*Nup107*^*KI/KI*^) or torsinA-KO (*Tor1a*^-/-^;*Nup107*^*KI/KI*^) mice were labeled with JFX554 HaloTag ligand overnight from DIV3 to DIV4 to saturate labeling of all existing Nup107. Following dye washout, some neurons were fixed and the rest were grown in JF646-containing media to label NPCs formed between DIV4 and time of fixation (Figure 4A). This pulse-chase labeling approach allowed us to distinguish NPCs formed up to DIV4 from those formed at DIV4-6, DIV4-8, or DIV4-10. We used this approach to test two possible mechanisms of NPC cluster formation: (i) rearrangement of NPCs post-formation, and (ii) improper localization of *de novo* assembling NPCs (Figure 4B). If NPCs rearrange into clusters post-formation in torsinA-KO neurons, clustering of both pulse- and chase-labeled NPCs is expected (Figure 4B, top panel). In contrast, if sites of new NPC biogenesis are mislocalized in the absence of torsinA, clustering of exclusively chase-labeled NPCs would be observed (Figure 4B, bottom panel).

**Figure 4:**
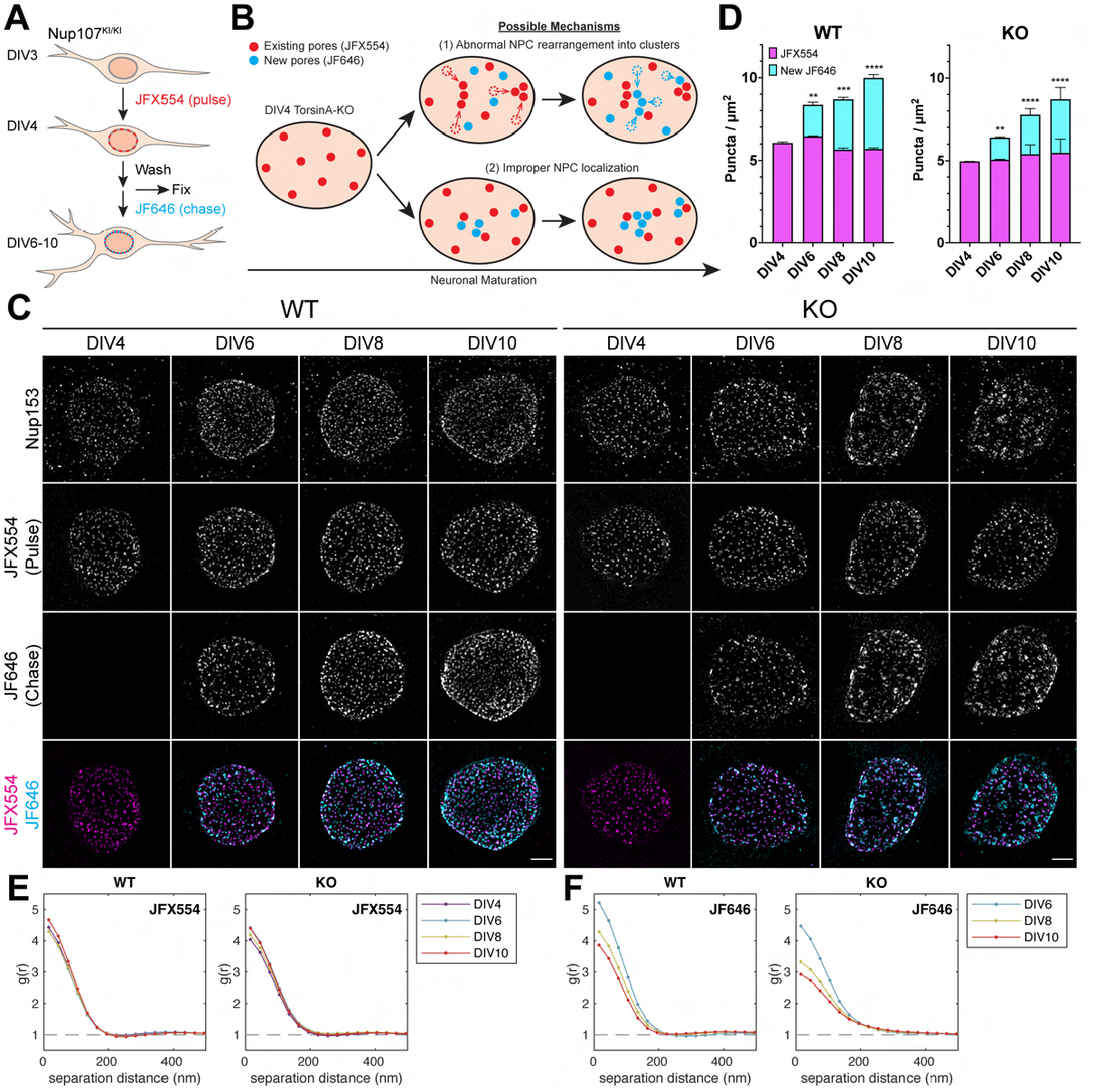
Sites of NPC biogenesis are abnormal in torsinA-KO neurons. **A**. Schematic of HaloTag pulse-chase experiment. **B**. Diagram of potential mechanisms for NPC clustering in torsinA-KO neurons. In (1), NPCs redistribute after formation, leading to clusters of existing and newly formed NPCs. In (2), sites of NPC biogenesis are mislocalized and clusters exclusively contain new NPCs. Red circles represent existing NPCs (pulse; JFX554). Blue circles represent new NPCs (chase; JF646). **C**. HaloTag pulse-chase SIM images of WT and torsinA-KO neurons. Neurons were stained with anti-Nup153 antibody post-fixation to label the total NPC population. Scale bar = 2μm. **D**. Density of JFX554 puncta and new JF646 puncta in DIV 4, 6, 8, and 10 WT and torsinA-KO HaloTag-Nup107 neurons. Plots show mean ± SD. Using DIV4 values as a reference, repeated measures two-way ANOVA with Dunnett’s test was performed. **P<0.01, ***P<0.001, ****P<0.0001. **E**. Autocorrelation of JFX554 images over 0-500nm separation distance. Similar starting amplitudes reflect constant JFX554 density. **F**. Autocorrelation of JF646 images over 0-500nm separation distance. Decreasing starting amplitudes over neuronal maturation reflect increasing JF646 density. Broadening of the curve in torsinA-KO neurons indicates spatial correlation of NPCs over larger distances.

Consistent with our previous findings, NPC biogenesis was upregulated in both genotypes and torsinA-KO neurons exhibited clustered NPCs (Figure 4C, Figure S5B). We used automated image analysis to identify NPCs formed up to DIV4 and those formed subsequently. The density of pulse-labeled (JFX554) NPCs remained constant over time. In contrast, the density of newly formed NPCs (new JF646; JF646-positive NPCs that do not overlap with JFX554) increased in both WT and torsinA-KO neurons (Figure 4D, Figure S5C, S5D), indicating successful pulse-chase labeling. JFX554-labeled NPCs maintain a sparse, dispersed distribution at all timepoints in both genotypes, even in DIV10 torsinA-KO neurons that contain large NPC clusters as observed via Nup153 labeling (Figure 4E, Figure S5E). In WT neurons, NPCs that form after DIV4 localize in empty spaces devoid of previously formed NPCs, preserving an overall uniform pattern of NPC distribution. In contrast, new NPCs that form after DIV4 in torsinA-KO neurons abnormally form in close proximity to one another, creating increasingly large clusters (Figure 4F, Figure S5F). These data support our second proposed mechanism in which mislocalization of newly forming NPCs leads to the development of NPC clusters in torsin-KO neurons. These data suggest that torsinA is an essential factor in determining sites of *de novo* NPC biogenesis. These experiments also demonstrate that in developing neurons, the spatial organization of NPCs is determined by the site of NPC formation rather than by post-formation rearrangement.

### NE blebs form at sites of NPC biogenesis

Abnormal NE blebs form selectively in post-migratory neurons in the torsinA-KO nervous system^35,39^. To further probe the potential link between NE bleb formation and NPC biogenesis, we explored the temporal and spatial relationship between these processes. To determine if NE blebs in torsinA-KO primary neurons form concomitantly with NPC biogenesis, we examined the nuclear membrane of WT and torsinA-KO neurons at DIV4 and DIV10 using transmission electron microscopy. We observed NE blebs in torsinA-KO neurons at both DIV4 and DIV10 (Figure 5A). Significantly more cells exhibited NE blebs at DIV10 (Figure 5B) and bleb frequency per cell increased dramatically from DIV4 to DIV10 (Figure 5C). NPC density increases almost twofold and abnormal NPC clusters emerge during this same period (DIV4–10), demonstrating a temporal link between these processes.

**Figure 5:**
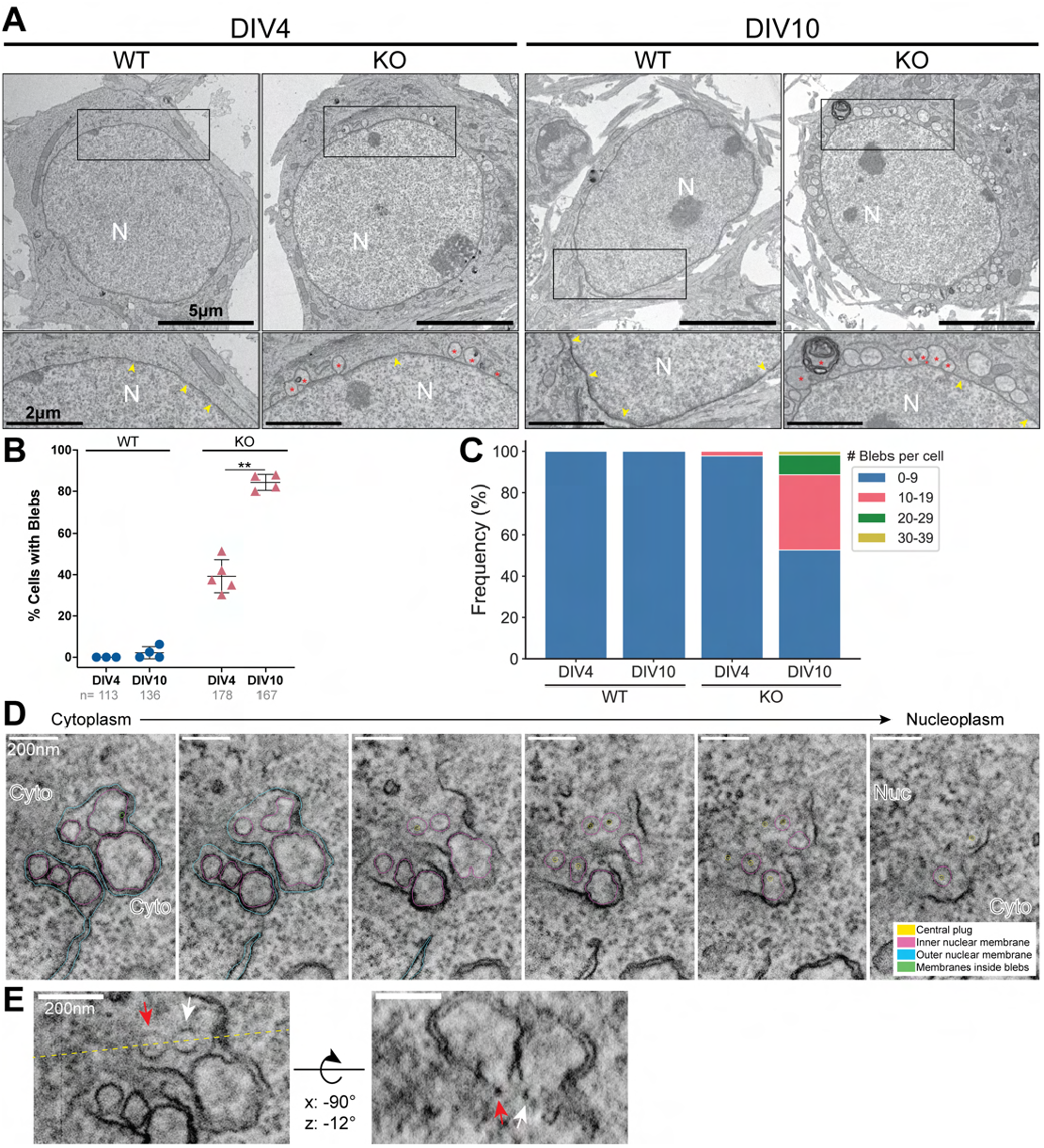
NE blebs spatially and temporally coincide with NPC biogenesis. **A**. Transmission electron microscopy (TEM) images of WT and torsinA-KO neurons at DIV4 and DIV10. N=nucleus. Fully-formed NPCs are marked with yellow arrowheads. NE blebs identified by blinded analysis are marked with red asterisks. Scale bar = 5μm for the top row, 2μm for inset. **B**. Quantitation of TEM images. Each point represents the % of cells with at least one NE bleb from each biological replicate. n shows the total number of cells analyzed across all replicates. **P=0.0043, paired t-test. **C**. Frequency of NE blebs per cell from all replicates. **D**. Slices from a DIV10 torsinA-KO tomogram overlayed with segmented contours of the outer nuclear membrane (cyan), inner nuclear membrane (magenta), membranes inside blebs (green), and central plug of pores (yellow). **E**. Rotated view of tomogram in (D) to show an XY view intersecting the center of two blebs at the yellow dashed line. Red and white arrows label pores with central plugs. The rotated view (right) demonstrates that these pores form the base of each bleb.

We next explored whether NE blebs and NPC clusters overlap spatially. One possibility is that areas of the NE with blebs are not suitable for NPC formation. Alternatively, colocalization of NPCs and NE blebs would support a mechanistic link between these processes. We tested these possibilities by generating 3D volumes of the WT and torsinA-KO DIV10 neuronal nuclear envelopes using intermediate high voltage scanning transmission electron microscopy (STEM) and multi-tilt electron tomography (Video 1). We observed NE blebs both as isolated entities and in clusters of several blebs enclosed within a single stretch of the outer nuclear membrane (Video 1, Figure S6A). In all cases, blebs connected to the nuclear membrane via pore channels (Figure 5D, 5E, Video 2) and we did not observe any NPC clusters between NE blebs. We did not observe any examples of vesicles reminiscent of large ribonucleoprotein granules reported in torsin mutant *Drosophila* cells^54^. Projection of all KO pores (regular and bleb-associated) clearly showed clustered distribution similar to SIM images of mature torsinA-KO neurons (Figure S6B). Considered together, the temporal and spatial connection between NE bleb and NPC formation support the hypothesis that NE blebs form at the sites of nascent NPCs in torsinA-KO neurons.

The 3D volumes also allowed us to analyze the dimensions of NPCs from WT and torsinA-KO neurons (Figure S6C). On average, the diameter of a pore in WT neurons was 80nm. Regular (not bleb-associated) pores in torsinA-KO neurons were slightly narrower, at 74.6nm, and bleb-associated KO pores were even smaller (69.8nm). These data are consistent with our dSTORM data and additionally demonstrate that even normal appearing NPCs in torsinA-KO neurons may not be able to dilate to the same degree as WT NPCs. We also found that the central plugs of bleb-associated pores facing the nucleoplasm are more electron dense compared to the plugs of regular pores, even within a single torsinA-KO neuron (Figure S6D). While the identity of these central plugs is unclear, they appear similar to plugs observed in immature NPCs^44,55^.

### Completion of NPC assembly is delayed in torsinA-KO neurons

NE blebs are transient in vivo, resolving without causing cell death^38,39^. We therefore explored whether blebs are also transient in torsinA-KO primary neurons. We examined the nuclear membrane of DIV10 and DIV18 WT and torsinA-KO neurons since our NPC density characterization suggests that NPC density may begin to plateau by DIV18. Both the prevalence and frequency of NE blebs decreased significantly between DIV10 and DIV18 in torsinA-KO neurons (Figure 6A-6C). In addition, we observed regions of the NE with multiple adjacent complete NPCs marked by fused INM and ONM in DIV18 torsinA-KO neurons (Figure 6A, last panel). We hypothesized that NPC assembly resumes and completes upon the resolution of NE blebs. To test this hypothesis and assess the NPC assembly state over time in torsinA-KO neurons, we labeled early- and late-recruited components, Nup153 and Nup358 respectively, at DIV10 and DIV18. At DIV10, Nup153 nuclear rim fluorescence intensity was similar between torsinA-KO and WT neurons, but Nup358 localization was reduced in torsinA-KO neurons (Figure 6D-6G). These data are consistent with our previous report^13^ and indicate that NPCs are stalled at an intermediate state in the absence of torsinA at this timepoint. At DIV18, however, Nup358 was successfully recruited to NPC clusters marked by Nup153 and the fluorescence intensity was restored (Figure 6G). We also validated this finding at single-NPC resolution using SIM (Figure S7A). Nup358 was sparsely distributed and failed to localize to clusters containing Nup153 in DIV10 torsinA-KO neurons. By DIV18, however, Nup358 exhibited clustered localization that overlapped with Nup153-positive clusters. Such a delay in NPC assembly completion was not observed in WT neurons, as most NPCs were labeled with both Nup153 and Nup358 at DIV10. Notably, NPCs remained clustered in DIV18 torsinA-KO neurons as observed by dSTORM and analyzed using autocorrelation (Figure S7B, S7C). These data indicate that torsinA is not required for fusion of the inner and outer nuclear membranes during interphase NPC assembly in neurons, as loss of torsinA causes delayed, rather than permanently halted, NPC biogenesis.

**Figure 6:**
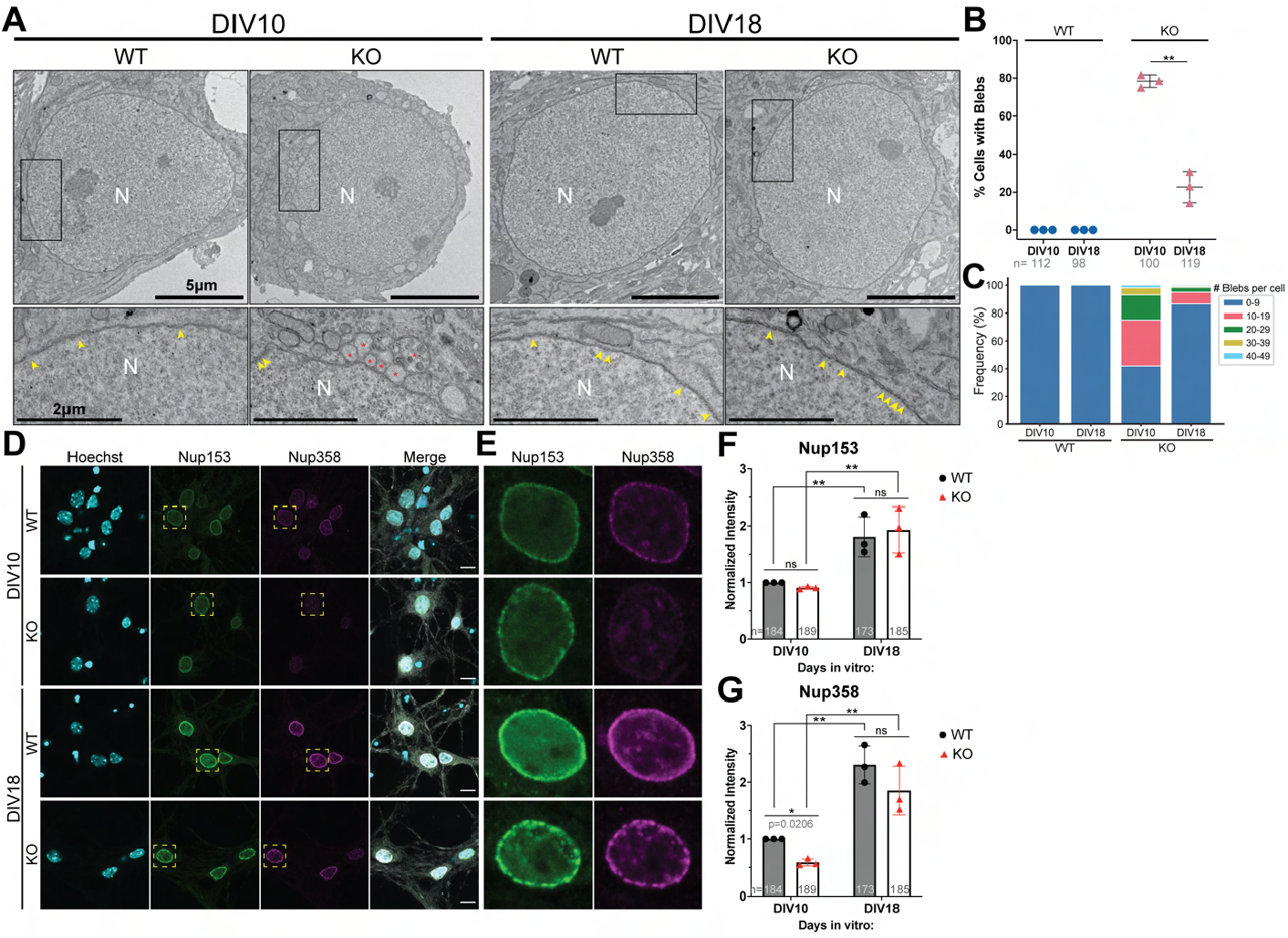
NPC assembly completes following NE bleb resolution. **A**. TEM images of WT and torsinA-KO neurons at DIV10 and DIV18. N=nucleus. Fully formed NPCs are marked with yellow arrowheads. NE blebs identified in blinded analysis are marked with red asterisks. Scale bar= 5μm for the top row, 2μm for inset. **B**. Quantitation of TEM images. Each point represents the % of cells with at least one NE bleb from each of 3 biological replicates. n shows the total number of cells that were analyzed across all replicates. **P=0.006; paired t-test. **C**. Frequency of NE blebs per cell from all replicates. **D**. Confocal images of DIV10 and DIV18 WT and torsinA-KO neurons labeled with Hoechst, Nup153, Nup358, and Map2 (shown in merge). Scale bar = 10μm. **E**. Zoomed in view of Nup153 and Nup358 channels of nuclei marked with yellow boxes in (D). **F, G**. Normalized Nup153 (F) and Nup358 (G) nuclear rim intensity of DIV10 and DIV18 WT and torsinA-KO primary neurons. ns, not significant; **P <0.01, repeated measures two-way ANOVA with Sidak’s multiple comparisons test.

## DISCUSSION

Our work establishes that torsinA is essential for the normal assembly kinetics and localization of interphase NPC biogenesis in neurons (Figure 7A, 7B). We find that NPC biogenesis is strongly and steadily upregulated during neuronal maturation, and that in torsinA-KO neurons, nascent NPCs localize abnormally into persisting, tightly packed clusters. In contrast to the aberrant spatial organization, the onset of NPC formation and total NPC density remain normal, suggesting that NPC number and NPC localization are governed by distinct mechanisms. We demonstrate directly using STEM tomography that inner nuclear membrane blebs in torsinA-KO neurons represent stalled intermediates of NPC assembly. Unlike the NPC clusters, however, these blebs resolve over time and allow NPC assembly to resume, resulting in delayed completion of NPC biogenesis. Considered together, our findings newly identify NPC upregulation as an important feature of neuronal maturation and provide insight into the poorly understood process of NPC spatial organization.

**Figure 7:**
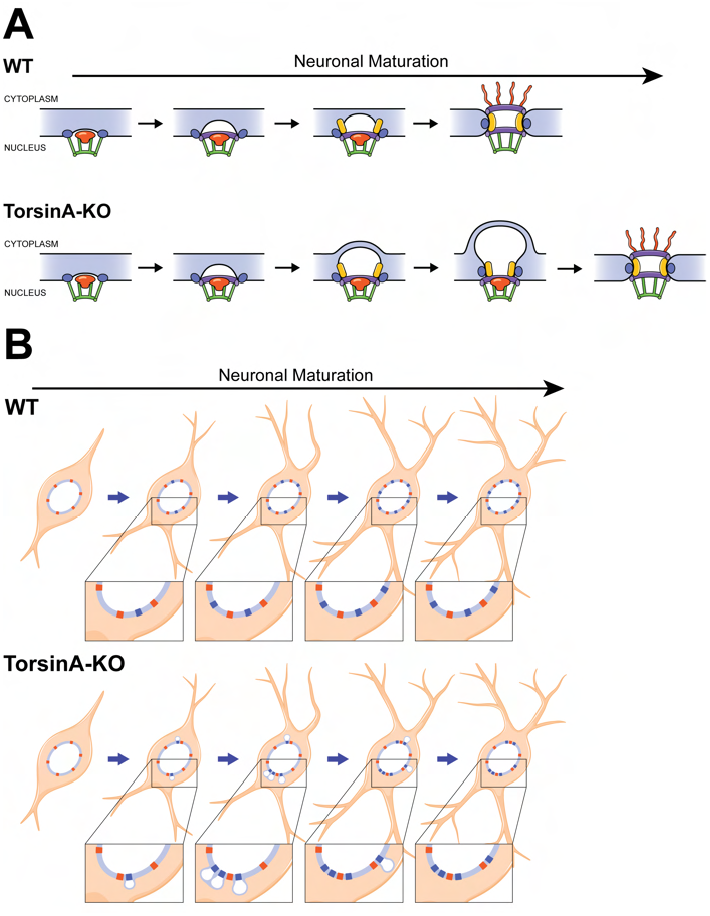
Summary of the effects of torsinA loss on NPC spatial organization and dynamics. **A**. Model of interphase NPC assembly in WT and torsinA-KO neurons. Onset of NPC assembly is not affected by torsinA deletion. Nuclear basket, inner ring, and transmembrane nucleoporins are recruited to the *de novo* NPC as the INM starts to bud. Instead of the normal INM-ONM fusion found in WT neurons, excessive INM extrusion causes NE blebs to emerge and enlarge in torsinA-KO neurons. These blebs stall torsinA-KO NPCs at an intermediate stage while WT NPC assembly completes. As torsinA-KO neurons continue to mature, NE blebs resolve and INM-ONM fusion occurs. Completion of NPC biogenesis is delayed in torsinA-KO neurons. **B**. Model of NPC localization in maturing WT (top) and torsinA-KO (bottom) neurons. In WT neurons, newly forming NPCs (blue) localize to empty spaces between existing NPCs (red), thereby maintaining uniform spatial organization. In maturing torsinA-KO neurons, newly forming NPCs (blue) localize abnormally close to each other or to existing NPCs (red), causing aberrant clusters. NPC biogenesis is upregulated in both genotypes and total NPC number is not affected by the absence of torsinA.

A tremendous challenge for post-migratory maturing neurons is meeting the large demand for protein synthesis, including mRNA transport, to supply the many proteins required to acquire characteristic neuronal phenotypes and to integrate into synaptically-connected circuits^48^. Considerable work in non-neuronal cells indicates that increasing NPC number is likely needed to support such metabolic demand^55–57^, but similar studies of NPC biogenesis have not been performed in neurons. To our knowledge, only one previous study has assessed NPC density during neural development, using freeze-fracture EM on brains from mice at four different ages^58^. Using multiple methods, including using super resolution microscopy with both antibody-dependent and -independent endogenous labeling of Nups, we demonstrate a doubling of NPC density from DIV4 to DIV18. This magnitude of increase is remarkably similar to the *in vivo* increase observed in the aforementioned EM study^58^. The robust twofold upregulation of neuronal NPC density contrasts with interphase NPC assembly in mitotic cells, which occurs concomitantly with NE expansion and results in little to no increase in NPC density^53,59–62^. While neuronal NPC biogenesis is sustained over a period of two weeks, interphase NPC assembly in mitotic cells completes on the scale of several hours^44,53,59–61,63^. Protein levels are upregulated during the first twenty days of maturing primary cortical neurons, coinciding with synaptogenesis and increasing complexity in neuronal morphology^48^. Neuronal NPC biogenesis likely occurs during a prolonged temporal window to accommodate an increased need for nucleocytoplasmic transport during neural development. Although NPC assembly in neurons and in interphasic mitotic cells both involve a *de novo* insertion of NPCs in intact nuclear membranes^44–46,60,61,64^, these differences in assembly timescale and the unique process of neuronal maturation raise the possibility of neuron-specific regulators of NPC biogenesis.

Our quantitative NPC distribution analyses demonstrate that WT NPCs are evenly dispersed at all timepoints assessed, suggesting the presence of a mechanism to maintain NPC spacing. Evenly dispersed NPCs have been observed in other cell types via EM studies^65^, but the mechanism that maintains NPC spatial organization is unknown. Our dSTORM studies clearly show that the nucleoporin clusters identified in confocal and SIM imaging contain NPCs with central pore channels, indicating a role for torsinA in neuronal NPC organization. Our data further suggest that independent pathways regulate NPC number and spatial organization: despite their mislocalization, the initiation of NPC assembly is unaffected as nucleoporins spanning multiple NPC subcomplexes are present in clusters (Nup153, Nup98, Nup210, and Nup107) and total NPC density is similar between WT and torsinA-KO neurons at all timepoints.

The pulse-chase studies of endogenous HaloTag-Nup107 demonstrate that torsinA loss disrupts the normal sites of NPC biogenesis. In maturing torsinA-KO neurons, NPCs form abnormally close to one another, implicating torsinA as a determinant of nascent NPC localization in neurons. We considered the possibility of NPC redistribution into clusters in the absence of torsinA since post-formation NPC rearrangement is observed in yeast^66,67^, and lamins, which are believed to anchor NPCs, appear abnormally shaped in some torsinA-KO neurons^13^. We find no evidence to support NPC redistribution into clusters; NPCs marked in immature torsinA-KO neurons are normally distributed and remain so throughout neuronal maturation. Most of these normally distributed NPCs are likely formed via a postmitotic NPC assembly mechanism prior to terminal neuronal differentiation, since torsinA loss does not cause NPC abnormalities in mitotic cells^40,42^. Indeed, interphase NPC assembly requires functional NPC transport^68–70^, and these normally distributed NPCs likely facilitate neuronal NPC biogenesis. Remarkably, these NPCs are not degraded during neuronal maturation as shown by our pulse-chase study, reflecting the long half-life of Nup107^52,53^ and lack of neuronal NPC turnover^19,20^.

How might torsinA ensure that nascent NPCs are properly localized? One possibility is that NPC formation requires a specific lipid microenvironment that is maintained by torsinA. The inner nuclear membrane (INM) is a metabolically active entity that can sense lipids^71–76^. The *Drosophila* torsin ortholog dTor maintains lipid homeostasis at the nuclear membrane^42,77,78^, and torsinA plays a role in mammalian hepatic lipid metabolism^79^. While no difference in lipids were found in mitotic cells lacking torsins^40^, NPC clustering is not observed in these nonneuronal cells. Future studies may reveal neural-specific spatial anomalies of specific lipids or lipid-binding proteins related to aberrant NPC localization. Another possibility is that torsinA may have a structural role in ensuring even distribution of INM proteins required for NPC assembly. Identification of torsinA interacting proteins in future studies will guide mechanistic studies on how torsinA participates in the proper localization of NPC biogenesis.

Our dSTORM and STEM data demonstrate that the central channel of the average NPC in DIV10 torsinA-KO neurons is slightly but significantly narrower than the average WT NPC. Interphase NPC assembly involves lateral dilation of the budding NPC prior to INM and ONM fusion^44–46^, suggesting that the smaller central channel of these NPCs may reflect stalled assembly intermediates that have not yet fully dilated. Consistent with this possibility, our EM data reveal that NE blebs emerge concurrently with upregulated NPC biogenesis and spatially correlate with mislocalized NPCs. These findings are similar to previous reports of NE blebs colocalizing with NPC components in mitotic cells lacking multiple torsin paralogs^40^. We additionally observe dense central plugs on the nucleoplasmic side of NE-bleb associated NPCs. These electron-dense plugs appear similar to plugs reported in newly forming pores^44,55^ and potentially represent unique machinery present in intermediate NPC structures.

Prior observations suggested that neuronal torsinA or torsins in mitotic cells may be involved in the INM-ONM fusion step of interphase NPC assembly. Chief among these observations is that Nup358, a cytoplasmic NPC component, is not recruited to Nup153-positive clusters in DIV10 torsinA-KO neurons^13^ or in interphase-formed NPCs in torsin-deficient mitotic cells^47^. Data from mitotic cells lacking torsins have also been used to support the hypothesis that NE blebs are a consequence of defective NPC assembly^47,80–82^. We demonstrate that torsinA is dispensable for INM-ONM fusion and that NPC biogenesis is delayed, rather than permanently halted, in the absence of torsinA. While DIV10 torsinA-KO neurons exhibit numerous NE blebs, these blebs resolve as the neurons mature, similar to the disappearance of blebs during CNS maturation *in vivo*^39^. NE bleb resolution is accompanied by the resumption of NPC assembly, as Nup358 is present in Nup153-positive NPC clusters in DIV18 torsinA-KO neurons. This contrasts sharply with findings in mitotic cells lacking multiple torsin paralogs^40,47,80^, in which completion of NPC assembly is not observed, possibly due to the difference in timescale of assembly between the cell types. Based on these new observations, we propose that NE blebs contribute to delayed NPC biogenesis by posing a physical barrier to the continuation of NPC assembly. The origin of NE blebs remains unclear. One possibility is that excessive INM budding occurs during NPC biogenesis in torsinA-KO neurons due to lipid environments unconducive to INM-ONM fusion. In this scenario, torsinA may play no direct role in NPC assembly, abnormalities of which may be a bystander effect of torsinA-related lipid defects^42,77,78^. NE blebs may resolve through correction of lipid composition via NE-ER junctions.

How might NPC abnormalities caused by torsinA deletion contribute to neural dysfunction? NPC function is compromised in maturing torsinA-KO neurons as demonstrated by abnormal protein nucleocytoplasmic transport^13^. Our STEM data indicate that even NPCs not associated with blebs in torsinA-KO neurons are smaller than WT NPCs, raising the intriguing possibility that full dilation of the NPC channel may be restricted regardless of assembly state. A persistently narrow channel size could limit the size of molecules that are able to diffuse through the NPCs. Such differences may negatively impact nucleocytoplasmic transport-dependent neural functions such as synaptic plasticity^3–6^, dysfunction of which is linked to the clinical manifestation of dystonia^83,84^. Delayed NPC assembly in maturing neurons may interfere with transport of locally translated RNA critical for neurodevelopmental processes such as dendritic outgrowth, axonal guidance and synaptogenesis^7–11^, which is altered in torsinA-KO^85^ and torsinA-ΔE knock-in brains^86–89^. Lastly, nucleoporins, including Nup153 and Nup98, physically interact with the genome and influence transcription^90–94^. NPC mislocalization may therefore lead to transcriptomic changes by altering the chromatin landscape.

TorsinA plays a unique and essential role during neural development. Neuronal torsinA levels are high during early postnatal development and decrease over subsequent weeks^39^. Deletion of torsinA from neural progenitors causes significant behavioral and neuropathological effects^95^, while eliminating torsinA in adult mice does not cause any phenotypes^43^. Conversely, restoring torsinA levels in juvenile, but not adult mice lacking torsinA, prevents such phenotypes, highlighting a critical window of torsinA function^43,96^. Our findings establish NPC biogenesis as a key molecular event of this neurodevelopmental critical period and raise the possibility that NPC-related processes may underlie additional critical periods in normal development as well as early-and late-onset neurological disorders.

## Acknowledgements

We thank the University of Michigan Microscopy Core facility for assistance with imaging and sample preparation. We thank D. Levin and X. Li for assistance with animal husbandry, A. Taylor for assistance with MATLAB code used for SIM analysis, S. Mosalaganti and A. Erwin for critical feedback on 3D-EM figures, D. Boassa for discussions about STEM, A. Moore for discussions about HaloTag ligands, A. Diehl for scientific illustrations, H. Worman and D. Yellajoshyula for critically reviewing the manuscript, and Dauer and Barmada laboratory members for additional discussions. This work was funded by support from NIH R01 NS110853, R01 DK118480, R01 NS077730 to W.T.D., NIH R01 NS097542, R01 NS113943, R56 NS128110 to S.J.B., NIH GM129347 and NSF MCB1552439 to S.L.V, and NIH T32 GM007315 and University of Michigan Rackham Predoctoral Fellowship to S.K. Electron tomography data acquisition, reconstruction and quantitative analyses of tomographic data were supported by NIH U24 NS120055, 1S10OD021784 and NSF2014862-UTA20-000890 to M.H.E. Deposition and management of acquired raw and derived electron tomography data within the Cell Image Library was further supported by NIH R01 GM082949 to M.H.E.

## Author contributions

S.K., S.S.P, S.J.B, and W.T.D conceived the study. S.K. performed all experiments and analyzed confocal and SIM data except autocorrelation, which S.L.V. analyzed. S.K. and S.L.V. acquired and processed dSTORM data and S.L.V. and T.R.S. conducted analyses. STEM tomograms were acquired and analyzed by S.P. in M.H.E’s laboratory. S.K. segmented the tomograms. S.L.V. prepared dSTORM and autocorrelation figures and S.K. prepared all other figures with input from S.S.P., S.J.B., and W.T.D. S.K. and S.S.P wrote the manuscript. S.L.V., S.J.B., and W.T.D. contributed to revisions. All authors reviewed the manuscript.

## METHODS

### Mice

All animal work was approved by the University of Michigan Institutional Animal Care and Use Committee. Experimental procedures were conducted in accordance with the approved protocols and the National Institutes of Health Guide for the care and use of laboratory animals. Mice were maintained at the University of Michigan in a temperature- and light-controlled animal facility room with access to food and water *ad libitum. Tor1a*^*-/-*^ and *Tor1a*^*+/+*^ mice were derived from *Tor1a*^*+/-*^ intercross^35^.

### Generation of HaloTag-Nup107 mouse line

The HaloTag-Nup107 mouse line was generated using CRISPR/Extreme Genome Editing technology in partnership with Biocytogen. HaloTag with a short linker and TEV site was inserted immediately upstream of the Nup107 start codon. *Nup107*^*KI/KI*^ mice were crossed into *Tor1a*^*+/-*^ line^35^ to generate *Nup107*^*KI/KI*^ *Tor1a*^*+/-*^ breeders. The following primers were used for genotyping: TGTGCTTCCGGAGAGCGGGAAG, ATAGGTTTGCCTGAAACTCCTGTGAC, and CGTGGTCGTCGAAGAAATAACCCAG. PCR cycle (95°C, 3 min; 95°C, 15 s; 70°C, 20 s (−0.7°C/c); 72°C, 1 min, 15 cycles; 98°C, 15 s; 60°C, 20 s; 72°C, 1 min, 25 cycles; 72°C, 7 min) yielded bands at 681bp for WT and 341bp for Nup107 knock-in.

### Primary neuron culturing

Brains were isolated from P0 pups into ice cold, filtered dissection buffer (6.85 mM sodium chloride, 0.27mM potassium chloride, 0.0085mM sodium phosphate dibasic anhydrous, 0.011mM potassium phosphate monobasic anhydrous, 33.3mM D-glucose, 43.8mM sucrose, 0.277mM HEPES, pH 7.4). After removing the cerebellum and the meninges, cortices were dissected out, placed into a microcentrifuge tube, and cut into small pieces with dissection forceps. Cortices were incubated in 50µL papain (2mg/mL; BrainBits) and 10µL DNase I (1mg/mL; Worthington Biochemical) for 30min at 37 °C. 500µL BrainPhys Neuronal Medium (Stemcell Technologies) and 10µL additional DNase I were added, and cortices were triturated using P1000 and P200 pipet tips. Triturated cortices were centrifuged at 1000rpm for 5min. After discarding the supernatants, the pellets were triturated and centrifuged three more times until the supernatant remained clear and neuronal pellets were visible. Pelleted neurons were resuspended in BrainPhys Neuronal medium with SM1 supplement, then plated onto 35mm #1.5 glass-bottom dishes (MatTek Life Sciences) or 12mm circular #1.5 coverslips (Neuvitro) coated with polyethylenimine (100 µg/ml; Polysciences). Neurons were incubated in 5% CO2 at 37 °C, with half media change every four days. Neurons derived from one animal were plated on several coverslips such that cells from a single biological replicate can be collected on different days for time course experiments.

### Immunofluorescence staining for confocal and structured illumination microscopy

Neurons plated on coverslips were rinsed with sterile Hank’s Balanced Salt Solution (HBSS) and fixed for 10min with pre-warmed 4% paraformaldehyde (PFA) (Electron Microscopy Sciences) in phosphate buffered saline (PBS). Fixed neurons were rinsed three times with PBS and permeabilized in 0.1% Triton X-100 (Millipore Sigma) in PBS for 10min. Neurons were then incubated in blocking buffer containing 5% bovine serium albumin (BSA) and 5% Normal Goat Serum (Jackson ImmunoResearch) for 30min, and labeled with primary antibodies diluted in PBS containing 0.1% normal goat serum overnight in 4 °C. The following day, neurons were washed three times in PBS, stained with secondary antibody for one hour, and washed three times with PBS. DNA was stained using Hoechst 33342 (1:10000 in PBS) for 5 minutes. Cells were washed twice with PBS and mounted in ProLong gold antifade mountant (ThermoFisher) for confocal or ProLong glass antifade mountant for SIM. For timecourse experiments, coverslips of neurons derived from the same animal were fixed on different days (timepoints). Coverslips fixed on earlier timepoints were stored in PBS at 4 °C until the end of the timecourse. Once all coverslips were collected, they were stained and mounted simultaneously.

### HaloTag-Nup107 labeling

For visualization of HaloTag-Nup107 in Fig. 3, neurons were incubated in media containing 50nM JF646 HaloTag ligand^97^ (Promega GA1120) or JFX554 HaloTag ligand^98^ (Kind gift of L. Lavis, Janelia) for 30 minutes. Excess ligands were then washed out by incubating labeled cells twice in neuronal media for >1h. Cells were then fixed and immunostained with antibodies. For pulse-chase labeling in Fig. 4, DIV3 neurons plated on glass coverslips were incubated in media containing 50nM JFX554 HaloTag ligand overnight. On DIV4, all coverslips were rinsed with HBSS, incubated in neuronal media twice for 1h each to wash out excess ligand, then rinsed again with HBSS. At least one coverslip was fixed to establish DIV4 NPC density. Remaining coverslips were incubated in conditioned neuronal media containing 50nM JF646 HaloTag ligand. Coverslips were subsequently washed and fixed on DIV6, 8, and 10. On DIV7, half of media was replaced with fresh conditioned media containing 50nM JF646. Conditioned neuronal media was generated by collecting media from excess neurons of the same age and combining 1:1 with fresh neuronal media. All rinses and media changes were done with care to ensure coverslips do not dry out. After DIV10, all fixed coverslips were immunostained with anti-Nup153 primary antibody and Alexafluor 488-conjugated secondary antibody to label total NPC population.

### dSTORM sample preparation and imaging

Neurons were rinsed with sterile HBSS and fixed for 10min with pre-warmed 4% PFA (Electron Microscopy Sciences) in PBS. Following fixation, neurons were rinsed three times with PBS and permeabilized in 0.2% Triton X-100 (Millipore Sigma) in PBS for 5min. Neurons were then incubated in blocking buffer containing 5% BSA for 30min, and labeled with Nup210 polyclonal antibody (Bethyl laboratories A301-795A; 1:200 in PBS) for two nights at 4 °C. Neurons were washed three times in PBS and stained with goat-anti-rabbit Alexa Fluor 647 Fab Fragment (1:800; Jackson ImmunoResearch 111-607-003) for an hour, washed three times with PBS, and imaged. Imaging and processing to determine localization and correct drift were conducted as previously described^99^. Samples were imaged in a buffer containing 100mM Tris, 25mM NaCl, 10% glucose, 1% β-mercaptoethanol, 500 μg/ml glucose oxidase (Sigma) and 80 μg/ml catalase (Sigma). Images were rendered by generating 2D histograms from localizations followed by convolution with a Gaussian for display purposes.

### Expansion microscopy

DIV10 neurons cultured on circular 12mm coverslips were fixed and immunostained with primary and secondary antibodies as described above. After the final wash following secondary antibody labeling, samples were incubated for 1h at room temperature in 25mM MA-NHS (methacrylic acid N-hydroxy succinimidyl ester) diluted in PBS^100^ then washed three times in PBS. Gelling chamber was assembled by covering a glass slide with a layer of parafilm and using stacks of two #1.5 square coverslips as spacers placed 8mm apart, in a setup similar to what was previously described^101^. Each coverslip was flipped onto spacers neuron-side down and the resulting chamber was gently filled with U-ExM gelling solution on ice^102^ (19% sodium acrylate, 10% acrylamide, 0.1% bis-acrylamide in PBS with 0.5% TEMED and 0.5% APS added last), then incubated in a humid container for 1h at 37 °C. Afterwards, gels was carefully isolated, digested with proteinase-K as in the U-ExM protocol, and allowed to expand for 30 minutes in a 6-well dish filled with ddH_2_O. Subsequently, gels were each moved to a 10cm dish, exchanged twice with fresh ddH_2_O, then allowed to expand overnight at 4 °C in fresh ddH_2_O. Approximately 4x expansion was estimated based on the width of the expanded gel. Fully expanded gels were cut in pieces and mounted cell-side-down on glass-bottomed 60mm dishes coated with poly-L-lysine. Gels were covered with clean ddH_2_O and imaged on ECLIPSE Ti inverted laser-scanning Nikon A1 confocal microscope with a 60x Apochromat water immersion (NA 1.2) objective using NIS-Elements AR (Nikon).

### Microscopy

SIM imaging was conducted on a Nikon N-SIM microscope equipped with an sCMOS camera (Flash 4.0; Hamamatsu Photonics, Japan) and LU-NV laser unit (Nikon) using a 100x Apochromat oil objective (NA 1.49). Z-stack acquisitions were centered around the flat plane of the nuclear envelope (NE) facing the adherent surface of the coverslip. 9 optical sections for dual-color imaging (Alexafluor488 and Alexafluor555 immunofluorescence labeling) and 7 optical sections for triple-color imaging (JFX554, JF646, and Alexafluor488 immunofluorescence labeling) were obtained at 200nm intervals. For reconstruction from raw data, illumination modulation contrast was set at 1 and high-resolution noise suppression was set between 0.1 and 0.8. After reconstruction, two optical slices most perfectly situated in the flat NE plane were combined into maximum intensity projection for analysis. For visualizing nucleoporin nuclear rim intensity, coverslips were imaged on ECLIPSE-Ti inverted laser-scanning Nikon A1 confocal microscope using an Apochromat Lambda 60x oil (NA 1.4, WD 0.13) objective. HaloTag-labeled neurons in Fig. 3b were imaged on an ECLIPSE-Ti2 Nikon inverted microscope equipped with a spinning-disk scan head (Yokogawa, CSU-X1) and an EM-CCD camera (iXon Ultra 888) with an Apochromat TIRF 60x oil objective (NA 1.49). All microscopes were operated with NIS-Elements AR (Nikon) and all imaging was conducted at room temperature. For presentation, images were adjusted for brightness and contrast. For experiments comparing fluorescence intensity, all adjustments were conducted using the same settings.

### Western blot

Brains from P0 mouse pups were isolated into ice cold, filtered dissection buffer as with primary neuron isolation. Dissected cortices were placed into a microcentrifuge tube containing 100μL lysis buffer (RIPA (ThermoFisher; #89900) with cOmplete Mini Protease Inhibitor Cocktail (Roche; #11836153001)) and homogenized using a plastic plunger followed by pipetting with a P200 pipet tip. Samples were spun down at 15,000 g and supernatants were collected. Pierce BCA protein assay (ThermoFisher; #23227) was performed to determine protein concentration and lysates were normalized to 1 μg/μL with lysis buffer. Normalized lysates were mixed with sample-loading buffer (Invitrogen NP0007) containing 2-mercaptoethanol (80μL added to 920μL sample-loading buffer) and boiled for 5 min at 95°C. 7.5 μg of protein and Precision Plus Kaleidoscope protein standard (Bio-Rad #1610375) were loaded and run on 4– 12% Bis-Tris gels (Invitrogen NP0323PK2). Protein was then transferred (350 mA for 1 hr at 4°C) onto a 0.22 μm PVDF membrane in transfer buffer containing 20% methanol.

Membranes were blocked in 5% non-fat dry milk in TBS-T (Tris-buffered saline (TBS) with 0.1% Tween-20), washed once in TBS-T, and cut horizontally at the 75kDa mark. Membranes were incubated in TBS-T containing primary antibodies overnight at 4°C. Top half was incubated in anti-Nup107 (1:1000; ab236634) and HaloTag (1:500; Promega G9211). Bottom half was incubated in anti-GAPDH (1:1000, MAB374) and anti-β-actin (1:1000, Cell Signaling 4967) The next day, membranes were washed three times with TBS-T, incubated 1h at room temperature in secondary antibodies, then imaged using Odyssey CLx. Donkey anti-mouse 680 and donkey anti-rabbit 800 IR secondaries were used at 1:10,000 dilution in 1:1 mixture of TBS-T and Intercept (TBS) Blocking buffer (LICOR 927-60000) containing 0.01% Tween-20.

### Transmission electron microscopy

Primary neurons were cultured on 35mm MatTek glass bottom dishes. For pre-fixation, cells were incubated in 1.25% glutaraldehyde in 0.05M cacodylate buffer at 37 °C for 5 minutes then 4°C overnight. Cells were post-fixed in a mixture of 1% osmium tetroxide (OsO_4_) and 1% potassium ferrocyanide [K4Fe(CN)_6_] in 0.1M cacodylate buffer. For improved contrast of the subcellular structures, the pre- and post-fixed cells were stained with 1% thiocarbohydrazide, additional 1% OsO4, 1% uranyl acetate, and Walton’s lead aspartate. Cells were dehydrated in ethanol, infiltrated in Spurr’s resin, and thermally polymerized at 70°C for 48 hours. A Leica EM UC7 ultramicrotome was used to cut 70nm ultrathin serial sections and the sections were placed on 300 mesh Cu bare grids. Sections were coated with 4nm carbon by a Leica EM ACE600 high vacuum coater, observed under a JEOL JEM-1400 Plus LaB6 transmission electron microscope at 60keV high tension, and imaged by an AMT NanoSprint 12 CMOS camera.

### Scanning transmission electron microscopy tomography

To reconstruct 3D representations of specimens, a high voltage transmission electron microscope (Titan Halo, 300kV; Thermo Fisher Scientific) was operated in scanning mode, enabling us to image thick sample sections (∼750nm). The sections were cut from a resin-embedded material processed for electron microscopy as described above, and then placed on 50nm Luxel film slot grids. After the grids were glow-discharged on both sides, gold particles of different size (10nm, 20nm and 50nm) were deposited on the sample surfaces for alignment purpose. The specimen under interest was tilted from −60° to +60° every 0.5° at four evenly distributed azimuthal angle positions as previously described^103^ and micrographs were collected on an annular dark field detector. Data acquisition was performed with SerialEM (version 3.6.12) at magnifications corresponding to pixel sizes in the range of 0.5nm to 2.5nm at full resolution. An iterative reconstruction procedure^103^ was used to reconstruct the final volumes. Segmentation and animations were generated manually using 3dmod software (version 4.11.7; University of Colorado Boulder, Boulder, CO, USA).

### STEM pore dimension analysis

To assess the dimensions of a pore assembly found in electron tomograms, the octagonal structure of the pore was segmented by placement of eight points on the nuclear side. Segmentation was done separately for each pore using the following steps: A small sub-volume containing the pore was first extracted from the full tomogram, and a preliminary segmentation of eight corner points was placed manually. Principal component analysis using the eight points was conducted to determine the octagon orientation, and the sub-volume was re-sliced to orient the pore in a front-facing view. The segmentation was then refined by readjusting the corner points and re-slicing the sub-volume. This process was iterated until a satisfactory segmentation of the pore was achieved. The diameter of each pore was calculated as *d / sin(*π*/8)* where *d* represents the average distance between two adjacent corner points of the segmented octagon. Segmentation was carried out with the 3dmod software.

### NPC segmentation from dSTORM

NPCs were segmented from 16 WT DIV10 neurons and 18 KO DIV10 neurons. Individual NPCs were segmented through a multi-step and fully automated process designed to identify objects of a particular size and aspect ratio while not biasing towards particles with distinct pores. First, an initial segmentation was conducted using a home-built implementation of DBSCAN^99,104^, with ε=40 nm and minPts = 25. The size of the major and minor axes of segmented objects were then determined by tabulating the eigenvalues of the covariance of x,y points with each segment. Single pores were defined as segments with major axes between 33 and 53nm, and minor axes between 23 and 43nm. For DBSCAN objects with major and minor axes larger than this range, we further segmented pores as follows: Images were reconstructed from points with 5nm bin sizes, then blurred with a disk-shaped filter with radii corresponding to 70nm. Local maxima were then identified in filtered images, then culled to remove maxima less than 90nm apart. Localizations acquired within 80nm of the remaining local maxima were associated as candidate particles. Candidate particles were retained if their major and minor axes met the same criteria as used in the initial DBSCAN segmentation.

Segmented pores were aligned using algorithms described previously^105^. To speed computation and estimate errors, localizations for a randomly chosen subset (250) of segmented pores were inputted into the alignment algorithm using 100nm pixels as units, with the “scale” parameter set to 10nm (0.1 in 100nm units). The final step was accomplished with imposed 50-fold symmetry to generate an angularly symmetric super particle. The density functions were tabulated by calculating the radial distance of super-particle localizations from the center of the particle, then binning to make a histogram, using bin-widths of Δr=2.5nm and each bin was divided by the total number of localizations in the super-particle, to correct for systematic differences in imaging conditions across conditions. Histograms were further divided by the area associated with each bin = 2πrΔr to obtain the normalized localization density at each radius. For each super-particle, the radius associated with the maximum localization density was estimated by fitting normalized histograms to a 2^nd^ order polynomial between 25 and 55nm, then returning the maximum of the fit. Significance between conditions was determined using a 2-tailed t-test of bootstrapped replicates through the ttest2 function in MATLAB.

Super-particles generated from WT DIV18 nuclei were used to validate the segmentation procedures used. NPCs in these nuclei can be densely distributed, leading to some particles segmented as large clusters in the initial DBSCAN segmentation.

### Autocorrelation analysis of dSTORM and SIM data

For dSTORM images, pair auto-correlations were tabulated from localizations within a user-defined mask, as described previously^99^. For SIM images, pair auto-correlations were tabulated from images with a user-defined mask, as described previously^106^. In some panels, curves are normalized to the value of the first spatial bin (corresponding to r<30nm) as normalized g(r) = 1+(g(r)-1)/(g(r<30nm)-1). In all cases, curves shown represent the average over cells with errors indicating the SEM over cell measurements.

### Calculation of nearest neighbor distances from dSTORM segmented NPCs

The centroids from each segmented NPC were determined by averaging the x and y coordinates of all localizations in the segmented NPC particle. Nearest neighbor distances were determined by tabulating pairwise distances between all NPC centroids with a cutoff of 1μm using the crosspairs_indicies() function in https://github.com/VeatchLab/smlm-analysis, then extracting the shortest distance associated with each NPC.

### Determination of nucleoporin puncta localization and density from SIM images

Nuclear ROIs were traced manually in FIJI^107^. For each image, a binary mask of ROIs was generated and saved. For images with high background signal (i.e. Nup antibody-labeled images), min/max histograms were adjusted on FIJI such that Otsu thresholding excluded nonspecific background signal. MATLAB scripts were used to obtain coordinates of nucleoporin puncta. First, all masks were eroded by 15px to exclude the nuclear rim where the nuclear envelope starts to curve, then overlayed with corresponding images after Otsu-thresholding. Then, imextendedmax (extended-maxima transform) was used on individual nuclei from these images to identify peaks of nucleoporin puncta. The x-y coordinate of each peak was recorded. If the peak of a puncta spanned two or more pixels, the centroid was used as the coordinate. Puncta density was calculated by dividing the number of unique peaks by the area of the eroded nuclear ROI. Outputs from the thresholding and masking step and images of unique peaks were saved as .tif files to enable visual inspection of proper peak identification.

### Nearest neighbor distance analysis from SIM images

The nearest neighbor distance between nucleoporin puncta and the number of puncta within a specified radius were calculated in MATLAB using the knnsearch() and rangesearch() functions, respectively.

### Nucleoporin puncta colocalization

Co-lococalization of puncta was determined based on assessment of whether peaks from one channel (“query”) overlay with the thresholded puncta of the other channel (“search”) using a custom written function. To determine % colocalization between Nup153 and HaloTag-Nup107 (Fig. 3), an image of unique peaks from the “query” channel were overlayed with the thresholded image of the “search” channel and the % of peaks whose coordinate’s pixel value in the “search” channel image is >0 was calculated. To determine the density and coordinates of new JF646 puncta in the HaloTag pulse-chase experiment (Fig. 4), this method was adapted such that JF646 peaks that did not colocalize with JFX554 puncta were counted.

### Nuclear rim fluorescence intensity measurement of Nups

Each channel of confocal z-stack was exported as a separate .tif file. Only Map2-positive neurons were analyzed. For each nucleus, the z-plane that most closely represented the middle plane of the nucleus was manually determined and an ROI of the nuclear rim was manually traced in FIJI. Using a custom-written script, ROIs was converted into 4px-wide bands containing only the nuclear periphery, which were then used to measure the fluorescence intensity of nucleoporins at the nuclear rim. For visualization, fluorescence intensity of each cell was normalized to the average intensity of the reference timepoint. For statistics, log of all raw intensity values were calculated to preserve the variability in the dataset, which is lost with normalization. Average log(intensity) values were calculated for biological replicates and used to compare timepoints.

### NE bleb analysis

TEM images were acquired such that one entire nucleus and associated NE blebs (if applicable) were visible per image. Upon imaging all coverslips, all folders and files were blinded such that genotype and neuronal age identifiers were removed. All nuclei were analyzed for 1) presence of blebs and 2) number of blebs. Number of blebs was determined by counting structures that fit the following criteria: 1) a clear bleb “neck” that connects to the inner nuclear membrane and is enveloped by the outer nuclear membrane, or 2) a membrane-bound vesicle-like structure directly adjacent to the nuclear membrane that is enveloped by the outer nuclear membrane. Note that this method of blinded quantification consistently undercounts NE blebs by excluding all structures that appear like double-membrane vesicles, which are visible portions of blebs whose connection to the inner nuclear membrane occurs on a different plane from the imaging plane. Following quantification, images were unblinded and grouped by biological replicate for processing.

### Data analysis and statistics

Statistical information, including n (total cells analyzed), mean, and statistical significance values, is indicated in the figures or figure legends. Data are plotted using superplots^108^ when applicable. N (number of biological replicates) is represented in superplots and statistical tests were performed on means of biological replicates. At least three biological replicates were used per experiment. Statistical tests were determined upon consultation with CSCAR (Consulting for Statistics, Computing, and Analytics Research) at the University of Michigan. Sphericity was assumed for ANOVA tests based on the Mauchly’s test of sphericity, which failed to reject the null hypothesis. Statistical significance was determined with Graphpad Prism 9 using the tests indicated in each figure. Data were considered statistically significant at ∗P < 0.05, ∗∗P < 0.01, ∗∗∗P < 0.001 and ∗∗∗∗P < 0.0001.

## Data and code availability

STEM volumes will be available on the Cell Image Library. FIJI, MATLAB, and python scripts for SIM and confocal analyses, as well as supplemental video files, are available on https://github.com/suminkim/NPC_NE_measurements.

## REFERENCES

1. Knockenhauer, K.E., and Schwartz, T.U. (2016). The Nuclear Pore Complex as a Flexible and Dynamic Gate. Cell 164, 1162–1171. 10.1016/j.cell.2016.01.034.

2. Beck, M., and Hurt, E. (2016). The nuclear pore complex: understanding its function through structural insight. Nat. Rev. Mol. Cell Biol. 18, 73–89. 10.1038/nrm.2016.147.

3. Kandel, E.R. (2001). The Molecular Biology of Memory Storage: A Dialogue Between Genes and Synapses. Science 294, 1030–1038. 10.1126/science.1067020.

4. Cohen, S., and Greenberg, M.E. (2008). Communication Between the Synapse and the Nucleus in Neuronal Development, Plasticity, and Disease. Annu. Rev. Cell Dev. Biol. 24, 183–209. 10.1146/annurev.cellbio.24.110707.175235.

5. Karpova, A., Mikhaylova, M., Bera, S., Bär, J., Reddy, P.P., Behnisch, T., Rankovic, V., Spilker, C., Bethge, P., Sahin, J., et al. (2013). Encoding and Transducing the Synaptic or Extrasynaptic Origin of NMDA Receptor Signals to the Nucleus. Cell 152, 1119–1133. 10.1016/j.cell.2013.02.002.

6. Kaushik, R., Grochowska, K.M., Butnaru, I., and Kreutz, M.R. (2014). Protein trafficking from synapse to nucleus in control of activity-dependent gene expression. Neuroscience 280, 340–350. 10.1016/j.neuroscience.2014.09.011.

7. Sutton, M.A., and Schuman, E.M. (2005). Local translational control in dendrites and its role in long-term synaptic plasticity. J. Neurobiol. 64, 116–131. 10.1002/neu.20152.

8. Sutton, M.A., and Schuman, E.M. (2006). Dendritic Protein Synthesis, Synaptic Plasticity, and Memory. Cell 127, 49–58. 10.1016/j.cell.2006.09.014.

9. Jung, H., Gkogkas, C.G., Sonenberg, N., and Holt, C.E. (2014). Remote Control of Gene Function by Local Translation. Cell 157, 26–40. 10.1016/j.cell.2014.03.005.

10. Batista, A.F.R., and Hengst, U. (2016). Intra-axonal protein synthesis in development and beyond. Int. J. Dev. Neurosci. 55, 140–149. 10.1016/j.ijdevneu.2016.03.004.

11. Holt, C.E., Martin, K.C., and Schuman, E.M. (2019). Local translation in neurons: visualization and function. Nat. Struct. Mol. Biol. 26, 557–566. 10.1038/s41594-019-0263-5.

12. Grima, J.C., Daigle, J.G., Arbez, N., Cunningham, K.C., Zhang, K., Ochaba, J., Geater, C., Morozko, E., Stocksdale, J., Glatzer, J.C., et al. (2017). Mutant Huntingtin Disrupts the Nuclear Pore Complex. Neuron. 10.1016/j.neuron.2017.03.023.

13. Pappas, S.S., Liang, C.C., Kim, S., Rivera, C.A.O., and Dauer, W.T. (2018). TorsinA dysfunction causes persistent neuronal nuclear pore defects. Hum. Mol. Genet. 27, 407–420. 10.1093/hmg/ddx405.

14. Eftekharzadeh, B., Daigle, J.G., Kapinos, L.E., Coyne, A., Schiantarelli, J., Carlomagno, Y., Cook, C., Miller, S.J., Dujardin, S., Amaral, A.S., et al. (2018). Tau Protein Disrupts Nucleocytoplasmic Transport in Alzheimer’s Disease. Neuron. 10.1016/j.neuron.2018.07.039.

15. Coyne, A.N., Zaepfel, B.L., Hayes, L., Fitchman, B., Salzberg, Y., Luo, E.-C., Bowen, K., Trost, H., Aigner, S., Rigo, F., et al. (2020). G4C2 Repeat RNA Initiates a POM121-Mediated Reduction in Specific Nucleoporins in C9orf72 ALS/FTD. Neuron 107, 1124–1140.e11. 10.1016/j.neuron.2020.06.027.

16. Lin, Y.-C., Kumar, M.S., Ramesh, N., Anderson, E.N., Nguyen, A.T., Kim, B., Cheung, S., McDonough, J.A., Skarnes, W.C., Lopez-Gonzalez, R., et al. (2021). Interactions between ALS-linked FUS and nucleoporins are associated with defects in the nucleocytoplasmic transport pathway. Nat. Neurosci. 24, 1077–1088. 10.1038/s41593-021-00859-9.

17. Basel-Vanagaite, L., Muncher, L., Straussberg, R., Pasmanik-Chor, M., Yahav, M., Rainshtein, L., Walsh, C.A., Magal, N., Taub, E., Drasinover, V., et al. (2006). Mutated nup62 causes autosomal recessive infantile bilateral striatal necrosis. Ann. Neurol. 60, 214–222. 10.1002/ana.20902.

18. Harrer, P., Schalk, A., Shimura, M., Baer, S., Calmels, N., Spitz, M.A., Warde, M.-T.A., Schaefer, E., Kittke, V.M.S., Dincer, Y., et al. (2023). Recessive NUP54 Variants Underlie Early-Onset Dystonia with Striatal Lesions. Ann. Neurol. 93, 330–335. 10.1002/ana.26544.

19. Savas, J.N., Toyama, B.H., Xu, T., Yates, J.R., and Hetzer, M.W. (2012). Extremely Long-Lived Nuclear Pore Proteins in the Rat Brain. Science 335, 942–942. 10.1126/science.1217421.

20. Toyama, B.H., Savas, J.N., Park, S.K., Harris, M.S., Ingolia, N.T., Yates, J.R., and Hetzer, M.W. (2013). Identification of Long-Lived Proteins Reveals Exceptional Stability of Essential Cellular Structures. Cell 154, 971–982. 10.1016/j.cell.2013.07.037.

21. Ozelius, L.J., Hewett, J.W., Page, C.E., Bressman, S.B., Kramer, P.L., Shalish, C., de Leon, D., Brin, M.F., Raymond, D., Corey, D.P., et al. (1997). The earlyonset torsion dystonia gene (DYT1) encodes an ATP-binding protein. Nat. Genet. 17, 40–48. 10.1038/ng0997-40.

22. Kustedjo, K., Bracey, M.H., and Cravatt, B.F. (2000). Torsin A and Its Torsion Dystonia-associated Mutant Forms Are Lumenal Glycoproteins That Exhibit Distinct Subcellular Localizations*. J. Biol. Chem. 275, 27933– 27939. 10.1074/jbc.M910025199.

23. Hewett, J., Ziefer, P., Bergeron, D., Naismith, T., Boston, H., Slater, D., Wilbur, J., Schuback, D., Kamm, C., Smith, N., et al. (2003). TorsinA in PC12 cells: Localization in the endoplasmic reticulum and response to stress. J. Neurosci. Res. 72, 158–168. 10.1002/jnr.10567.

24. Goodchild, R.E., and Dauer, W.T. (2004). Mislocalization to the nuclear envelope: An effect of the dystonia-causing torsinA mutation. Proc. Natl. Acad. Sci. 101, 847–852. 10.1073/pnas.0304375101.

25. Naismith, T.V., Heuser, J.E., Breakefield, X.O., and Hanson, P.I. (2004). TorsinA in the nuclear envelope. Proc. Natl. Acad. Sci. 101, 7612–7617. 10.1073/pnas.0308760101.

26. Gonzalez-Alegre, P., and Paulson, H.L. (2004). Aberrant Cellular Behavior of Mutant TorsinA Implicates Nuclear Envelope Dysfunction in DYT1 Dystonia. J. Neurosci. 24, 2593–2601. 10.1523/JNEUROSCI.4461-03.2004.

27. Gerace, L. (2004). TorsinA and torsion dystonia: Unraveling the architecture of the nuclear envelope. Proc. Natl. Acad. Sci. 101, 8839–8840. 10.1073/pnas.0402441101.

28. Goodchild, R.E., and Dauer, W.T. (2005). The AAA+ protein torsinA interacts with a conserved domain present in LAP1 and a novel ER protein. J. Cell Biol. 168, 855–862. 10.1083/jcb.200411026.

29. Callan, A.C., Bunning, S., Jones, O.T., High, S., and Swanton, E. (2006). Biosynthesis of the dystonia-associated AAA+ ATPase torsinA at the endoplasmic reticulum. Biochem. J. 401, 607–612. 10.1042/BJ20061313.

30. Ozelius, L.J., Hewett, J., Kramer, P., Bressman, S.B., Shalish, C., De Leon, D., Rutter, M., Risch, N., Brin, M.F., Markova, E.D., et al. (1997). Fine localization of the torsion dystonia gene (DYT1) on human chromosome 9q34: YAC map and linkage disequilibrium. Genome Res. 7, 483–494. 10.1101/gr.7.5.483.

31. Ozelius, L.J., Page, C.E., Klein, C., Hewett, J.W., Mineta, M., Leung, J., Shalish, C., Bressman, S.B., de Leon, D., Brin, M.F., et al. (1999). The TOR1A (DYT1) gene family and its role in early onset torsion dystonia. Genomics 62, 377–384. 10.1006/geno.1999.6039.

32. Nery, F.C., Zeng, J., Niland, B.P., Hewett, J., Farley, J., Irimia, D., Li, Y., Wiche, G., Sonnenberg, A., and Breakefield, X.O. (2008). TorsinA binds the KASH domain of nesprins and participates in linkage between nuclear envelope and cytoskeleton. J. Cell Sci. 121, 3476–3486. 10.1242/jcs.029454.

33. Vander Heyden, A.B., Naismith, T.V., Snapp, E.L., Hodzic, D., and Hanson, P.I. (2009). LULL1 retargets TorsinA to the nuclear envelope revealing an activity that is impaired by the DYT1 dystonia mutation. Mol. Biol. Cell 20, 2661–2672. 10.1091/mbc.E09-01-0094.

34. VanGompel, M.J.W., Nguyen, K.C.Q., Hall, D.H., Dauer, W.T., and Rose, L.S. (2015). A novel function for the Caenorhabditis elegans torsin OOC-5 in nucleoporin localization and nuclear import. Mol. Biol. Cell 26, 1752–1763. 10.1091/mbc.E14-07-1239.

35. Goodchild, R.E., Kim, C.E., and Dauer, W.T. (2005). Loss of the dystonia-associated protein torsinA selectively disrupts the neuronal nuclear envelope. Neuron 48, 923–932. 10.1016/j.neuron.2005.11.010.

36. Zhao, C., Brown, R.S.H., Chase, A.R., Eisele, M.R., and Schlieker, C. (2013). Regulation of Torsin ATPases by LAP1 and LULL1. Proc. Natl. Acad. Sci. 110, E1545– E1554. 10.1073/pnas.1300676110.

37. Rose, A.E., Brown, R.S.H., and Schlieker, C. (2015). Torsins: not your typical AAA+ ATPases. Crit. Rev. Biochem. Mol. Biol. 50, 532–549. 10.3109/10409238.2015.1091804.

38. Pappas, S.S., Darr, K., Holley, S.M., Cepeda, C., Mabrouk, O.S., Wong, J.M.T., LeWitt, T.M., Paudel, R., Houlden, H., Kennedy, R.T., et al. (2015). Forebrain deletion of the dystonia protein torsinA causes dystonic-like movements and loss of striatal cholinergic neurons. eLife 4, e08352. 10.7554/eLife.08352.

39. Tanabe, L.M., Liang, C.C., and Dauer, W.T. (2016). Neuronal Nuclear Membrane Budding Occurs during a Developmental Window Modulated by Torsin Paralogs. Cell Rep. 16, 3322–3333. 10.1016/j.celrep.2016.08.044.

40. Laudermilch, E., Tsai, P.-L., Graham, M., Turner, E., Zhao, C., and Schlieker, C. (2016). Dissecting Torsin/cofactor function at the nuclear envelope: a genetic study. Mol. Biol. Cell 27, 3964–3971. 10.1091/mbc.E16-07-0511.

41. Kim, C.E., Perez, A., Perkins, G., Ellisman, M.H., and Dauer, W.T. (2010). A molecular mechanism underlying the neural-specific defect in torsinA mutant mice. Proc. Natl. Acad. Sci. 107, 9861–9866. 10.1073/pnas.0912877107.

42. Jacquemyn, J., Foroozandeh, J., Vints, K., Swerts, J., Verstreken, P., Gounko, N.V., Gallego, S.F., and Goodchild, R. (2021). Torsin and NEP1R1-CTDNEP1 phosphatase affect interphase nuclear pore complex insertion by lipid-dependent and lipid-independent mechanisms. EMBO J. 40, e106914. 10.15252/embj.2020106914.

43. Li, J., Levin, D.S., Kim, A.J., Pappas, S.S., and Dauer, W.T. (2021). TorsinA restoration in a mouse model identifies a critical therapeutic window for DYT1 dystonia. J. Clin. Invest. 131. 10.1172/JCI139606.

44. Otsuka, S., Bui, K.H., Schorb, M., Hossain, M.J., Politi, A.Z., Koch, B., Eltsov, M., Beck, M., and Ellenberg, J. (2016). Nuclear pore assembly proceeds by an insideout extrusion of the nuclear envelope. eLife 5, e19071. 10.7554/eLife.19071.

45. Otsuka, S., and Ellenberg, J. (2018). Mechanisms of nuclear pore complex assembly – two different ways of building one molecular machine. FEBS Lett. 592, 475– 488. 10.1002/1873-3468.12905.

46. Otsuka, S., Tempkin, J.O.B., Zhang, W., Politi, A.Z., Rybina, A., Hossain, M.J., Kueblbeck, M., Callegari, A., Koch, B., Morero, N.R., et al. (2023). A quantitative map of nuclear pore assembly reveals two distinct mechanisms. Nature 613, 575–581. 10.1038/s41586-022-05528-w.

47. Rampello, A.J., Laudermilch, E., Vishnoi, N., Prophet, S.M., Shao, L., Zhao, C., Lusk, C.P., and Schlieker, C. (2020). Torsin ATPase deficiency leads to defects in nuclear pore biogenesis and sequestration of MLF2. J. Cell Biol. 219, e201910185. 10.1083/jcb.201910185.

48. Lesuisse, C., and Martin, L.J. (2002). Long-term culture of mouse cortical neurons as a model for neuronal development, aging, and death. J. Neurobiol. 51, 9–23. 10.1002/neu.10037.

49. Mosalaganti, S., Kosinski, J., Albert, S., Schaffer, M., Strenkert, D., Salomé, P.A., Merchant, S.S., Plitzko, J.M., Baumeister, W., Engel, B.D., et al. (2018). In situ architecture of the algal nuclear pore complex. Nat. Commun. 9, 2361. 10.1038/s41467-018-04739-y.

50. Lin, D.H., and Hoelz, A. (2019). The Structure of the Nuclear Pore Complex (An Update). Annu. Rev. Biochem. 88, 725–783. 10.1146/annurev-biochem-062917-011901.

51. Los, G.V., Encell, L.P., McDougall, M.G., Hartzell, D.D., Karassina, N., Zimprich, C., Wood, M.G., Learish, R., Ohana, R.F., Urh, M., et al. (2008). HaloTag: A Novel Protein Labeling Technology for Cell Imaging and Protein Analysis. ACS Publ. 10.1021/cb800025k.

52. Rabut, G., Doye, V., and Ellenberg, J. (2004). Mapping the dynamic organization of the nuclear pore complex inside single living cells. Nat. Cell Biol. 6, 1114–1121. 10.1038/ncb1184.

53. Dultz, E., and Ellenberg, J. (2010). Live imaging of single nuclear pores reveals unique assembly kinetics and mechanism in interphase. J. Cell Biol. 191, 15–22. 10.1083/jcb.201007076.

54. Jokhi, V., Ashley, J., Nunnari, J., Noma, A., Ito, N., Wakabayashi-Ito, N., Moore, M.J., and Budnik, V. (2013). Torsin Mediates Primary Envelopment of Large Ribonucleoprotein Granules at the Nuclear Envelope. Cell Rep. 3, 988–995. 10.1016/j.celrep.2013.03.015.

55. Maul, G.G., Price, J.W., and Lieberman, M.W. (1971). Formation and Distribution of Nuclear Pore Complexes in Interphase. J. Cell Biol. 51, 405–418. 10.1083/jcb.51.2.405.

56. Maul, G., and Deaven, L. (1977). Quantitative determination of nuclear pore complexes in cycling cells with differing DNA content. J. Cell Biol. 73, 748–760. 10.1083/jcb.73.3.748.

57. Maul, G.G., Deaven, L.L., Freed, J.J., Campbell, L.M., and Beçak, W. (1980). Investigation of the determinants of nuclear pore number. Cytogenet. Genome Res. 26, 175–190. 10.1159/000131439.

58. Lodin, Z., Blumajer, J., and Mares, V. (1978). Nuclear pore complexes in cells of the developing mouse cerebral cortex. Acta Histochem. 63, 74–79. 10.1016/S0065-1281(78)80009-9.

59. Maul, G.G., Maul, H.M., Scogna, J.E., Lieberman, M.W., Stein, G.S., Hsu, B.Y.-L., and Borun, T.W. (1972). Time Sequence of Nuclear Pore Formation in Phytohemagglutinin-stimulated Lymphocytes and in HeLa Cells during the Cell Cycle. J. Cell Biol. 55, 433– 447. 10.1083/jcb.55.2.433.

60. D’Angelo, M.A., Anderson, D.J., Richard, E., and Hetzer, M.W. (2006). Nuclear Pores Form de Novo from Both Sides of the Nuclear Envelope. Science 312, 440–443. 10.1126/science.1124196.

61. Maeshima, K., Iino, H., Hihara, S., Funakoshi, T., Watanabe, A., Nishimura, M., Nakatomi, R., Yahata, K., Imamoto, F., Hashikawa, T., et al. (2010). Nuclear pore formation but not nuclear growth is governed by cyclin-dependent kinases (Cdks) during interphase. Nat. Struct. Mol. Biol. 17, 1065–1071. 10.1038/nsmb.1878.

62. Varberg, J.M., Unruh, J.R., Bestul, A.J., Khan, A.A., and Jaspersen, S.L. (2022). Quantitative analysis of nuclear pore complex organization in Schizosaccharomyces pombe. Life Sci. Alliance 5. 10.26508/lsa.202201423.

63. Doucet, C.M., Talamas, J.A., and Hetzer, M.W. (2010). Cell Cycle-Dependent Differences in Nuclear Pore Complex Assembly in Metazoa. 10.1016/j.cell.2010.04.036.

64. Doucet, C.M., and Hetzer, M.W. (2010). Nuclear pore biogenesis into an intact nuclear envelope. Chromosoma 119, 469–477. 10.1007/s00412-010-0289-2.

65. Maul, G.G. (1977). The nuclear and the cytoplasmic pore complex: structure, dynamics, distribution, and evolution. Int. Rev. Cytol. Suppl., 75–186.

66. Bucci, M., and Wente, S.R. (1997). In vivo dynamics of nuclear pore complexes in yeast. J. Cell Biol. 136, 1185–1199. 10.1083/jcb.136.6.1185.

67. Belgareh, N., and Doye, V. (1997). Dynamics of Nuclear Pore Distribution in Nucleoporin Mutant Yeast Cells. J. Cell Biol. 136, 747–759.

68. Mattaj, I.W. (2004). Sorting out the nuclear envelope from the endoplasmic reticulum. Nat. Rev. Mol. Cell Biol. 5, 65–69. 10.1038/nrm1263.

69. Iino, H., Maeshima, K., Nakatomi, R., Kose, S., Hashikawa, T., Tachibana, T., and Imamoto, N. (2010). Live imaging system for visualizing nuclear pore complex (NPC) formation during interphase in mammalian cells. Genes Cells 15, 647–660. 10.1111/j.1365-2443.2010.01406.x.

70. Vollmer, B., Lorenz, M., Moreno-Andrés, D., Bodenhöfer, M., De Magistris, P., Astrinidis, S.A., Schooley, A., Flötenmeyer, M., Leptihn, S., and Antonin, W. (2015). Nup153 Recruits the Nup107-160 Complex to the Inner Nuclear Membrane for Interphasic Nuclear Pore Complex Assembly. Dev. Cell 33, 717–728. 10.1016/j.devcel.2015.04.027.

71. Romanauska, A., and Köhler, A. (2018). The Inner Nuclear Membrane Is a Metabolically Active Territory that Generates Nuclear Lipid Droplets. Cell 174, 700–715.e18. 10.1016/j.cell.2018.05.047.

72. Ohsaki, Y., Kawai, T., Yoshikawa, Y., Cheng, J., Jokitalo, E., and Fujimoto, T. (2016). PML isoform II plays a critical role in nuclear lipid droplet formation. J. Cell Biol. 212, 29–38. 10.1083/jcb.201507122.

73. Merta, H., and Bahmanyar, S. (2018). The Inner Nuclear Membrane Takes On Lipid Metabolism. Dev. Cell 47, 397–399. 10.1016/j.devcel.2018.11.005.

74. Sołtysik, K., Ohsaki, Y., Tatematsu, T., Cheng, J., and Fujimoto, T. (2019). Nuclear lipid droplets derive from a lipoprotein precursor and regulate phosphatidylcholine synthesis. Nat. Commun. 10, 473. 10.1038/s41467-019-08411-x.

75. Sołtysik, K., Ohsaki, Y., Tatematsu, T., Cheng, J., Maeda, A., Morita, S., and Fujimoto, T. (2021). Nuclear lipid droplets form in the inner nuclear membrane in a seipin-independent manner. J. Cell Biol. 220, e202005026. 10.1083/jcb.202005026.

76. Lee, S., Merta, H., Rodríguez, J.W.C., and Bahmanyar, S. (2022). A membrane sensing mechanism couples local lipid metabolism to protein degradation at the inner nuclear membrane. 2022.07.06.498903. 10.1101/2022.07.06.498903.

77. Grillet, M., Dominguez Gonzalez, B., Sicart, A., Pöttler, M., Cascalho, A., Billion, K., Hernandez Diaz, S., Swerts, J., Naismith, T.V., Gounko, N.V., et al. (2016). Torsins Are Essential Regulators of Cellular Lipid Metabolism. Dev. Cell 38, 235–247. 10.1016/j.devcel.2016.06.017.

78. Cascalho, A., Foroozandeh, J., Hennebel, L., Swerts, J., Klein, C., Rous, S., Dominguez Gonzalez, B., Pisani, A., Meringolo, M., Gallego, S.F., et al. (2020). Excess Lipin enzyme activity contributes to TOR1A recessive disease and DYT-TOR1A dystonia. Brain 143, 1746–1765. 10.1093/brain/awaa139.

79. Shin, J.-Y., Hernandez-Ono, A., Fedotova, T., Östlund, C., Lee, M.J., Gibeley, S.B., Liang, C.-C., Dauer, W.T., Ginsberg, H.N., and Worman, H.J. (2019). Nuclear envelope–localized torsinA-LAP1 complex regulates hepatic VLDL secretion and steatosis. J. Clin. Invest. 129, 4885–4900. 10.1172/JCI129769.

80. Prophet, S.M., Rampello, A.J., Niescier, R.F., Gentile, J.E., Mallik, S., Koleske, A.J., and Schlieker, C. (2022). Atypical nuclear envelope condensates linked to neurological disorders reveal nucleoporin-directed chaperone activities. Nat. Cell Biol. 24, 1630–1641. 10.1038/s41556-022-01001-y.

81. Kuiper, E.F.E., Gallardo, P., Bergsma, T., Mari, M., Kolbe Musskopf, M., Kuipers, J., Giepmans, B.N.G., Steen, A., Kampinga, H.H., Veenhoff, L.M., et al. (2022). The chaperone DNAJB6 surveils FG-nucleoporins and is required for interphase nuclear pore complex biogenesis. Nat. Cell Biol. 24, 1584– 1594. 10.1038/s41556-022-01010-x.

82. Keuenhof, K.S., Kohler, V., Broeskamp, F., Panagaki, D., Speese, S.D., Büttner, S., and Höög, J.L. (2023). Nuclear envelope budding and its cellular functions. Nucleus 14, 2178184. 10.1080/19491034.2023.2178184.

83. Quartarone, A., and Hallett, M. (2013). Emerging concepts in the physiological basis of dystonia. Mov. Disord. Off. J. Mov. Disord. Soc. 28, 958–967. 10.1002/mds.25532.

84. Quartarone, A., and Ghilardi, M.F. (2022). Chapter 14 - Neuroplasticity in dystonia: Motor symptoms and beyond. In Handbook of Clinical Neurology Neuroplasticity., A. Quartarone, M. F. Ghilardi, and F. Boller, eds. (Elsevier), pp. 207–218. 10.1016/B978-0-12-819410-2.00031-X.

85. Yellajoshyula, D., Opeyemi, S., Dauer, W.T., and Pappas, S.S. (2022). Genetic evidence of aberrant striatal synaptic maturation and secretory pathway alteration in a dystonia mouse model. Dystonia 0. 10.3389/dyst.2022.10892.

86. Song, C.-H., Bernhard, D., Bolarinwa, C., Hess, E.J., Smith, Y., and Jinnah, H.A. (2013). Subtle microstructural changes of the striatum in a DYT1 knock-in mouse model of dystonia. Neurobiol. Dis. 54, 362–371. 10.1016/j.nbd.2013.01.008.

87. Song, C.-H., Bernhard, D., Hess, E.J., and Jinnah, H.A. (2014). Subtle microstructural changes of the cerebellum in a knock-in mouse model of DYT1 dystonia. Neurobiol. Dis. 62, 372–380. 10.1016/j.nbd.2013.10.003.

88. Vanni, V., Puglisi, F., Bonsi, P., Ponterio, G., Maltese, M., Pisani, A., and Mandolesi, G. (2015). Cerebellar synaptogenesis is compromised in mouse models of DYT1 dystonia. Exp. Neurol. 271, 457–467. 10.1016/j.expneurol.2015.07.005.

89. Maltese, M., Stanic, J., Tassone, A., Sciamanna, G., Ponterio, G., Vanni, V., Martella, G., Imbriani, P., Bonsi, P., Mercuri, N.B., et al. (2018). Early structural and functional plasticity alterations in a susceptibility period of DYT1 dystonia mouse striatum. eLife 7, e33331. 10.7554/eLife.33331.

90. Capelson, M., Liang, Y., Schulte, R., Mair, W., Wagner, U., and Hetzer, M.W. (2010). Chromatin-Bound Nuclear Pore Components Regulate Gene Expression in Higher Eukaryotes. Cell 140, 372–383. 10.1016/j.cell.2009.12.054.

91. Ibarra, A., and Hetzer, M.W. (2015). Nuclear pore proteins and the control of genome functions. Genes Dev. 29, 337–349. 10.1101/gad.256495.114.

92. Pascual-Garcia, P., Debo, B., Aleman, J.R., Talamas, J.A., Lan, Y., Nguyen, N.H., Won, K.J., and Capelson, M. (2017). Metazoan Nuclear Pores Provide a Scaffold for Poised Genes and Mediate Induced Enhancer-Promoter Contacts. Mol. Cell 66, 63–76.e6. 10.1016/j.molcel.2017.02.020.

93. Sun, J., Shi, Y., and Yildirim, E. (2019). The Nuclear Pore Complex in Cell Type-Specific Chromatin Structure and Gene Regulation. Trends Genet. 35, 579–588. 10.1016/j.tig.2019.05.006.

94. Kadota, S., Ou, J., Shi, Y., Lee, J.T., Sun, J., and Yildirim, E. (2020). Nucleoporin 153 links nuclear pore complex to chromatin architecture by mediating CTCF and cohesin binding. Nat. Commun. 11, 2606. 10.1038/s41467-020-16394-3.

95. Liang, C.C., Tanabe, L.M., Jou, S., Chi, F., and Dauer, W.T. (2014). TorsinA hypofunction causes abnormal twisting movements and sensorimotor circuit neurodegeneration. J. Clin. Invest. 124, 3080–3092. 10.1172/JCI72830.

96. Li, J., Kim, S., Pappas, S.S., and Dauer, W.T. (2021). CNS critical periods: implications for dystonia and other neurodevelopmental disorders. JCI Insight 6. 10.1172/jci.insight.142483.

97. Grimm, J.B., English, B.P., Chen, J., Slaughter, J.P., Zhang, Z., Revyakin, A., Patel, R., Macklin, J.J., Normanno, D., Singer, R.H., et al. (2015). A general method to improve fluorophores for live-cell and singlemolecule microscopy. Nat. Methods 12, 244–250. 10.1038/nmeth.3256.

98. Grimm, J.B., Xie, L., Casler, J.C., Patel, R., Tkachuk, A.N., Falco, N., Choi, H., Lippincott-Schwartz, J., Brown, T.A., Glick, B.S., et al. (2021). A General Method to Improve Fluorophores Using Deuterated Auxochromes. JACS Au 1, 690–696. 10.1021/jacsau.1c00006.

99. Shaw, T.R., Fazekas, F.J., Kim, S., Flanagan-Natoli, J.C., Sumrall, E.R., and Veatch, S.L. (2022). Estimating the localization spread function of static single-molecule localization microscopy images. Biophys. J. 121, 2906– 2920. 10.1016/j.bpj.2022.06.036.

100. Chozinski, T.J., Halpern, A.R., Okawa, H., Kim, H.-J., Tremel, G.J., Wong, R.O.L., and Vaughan, J.C. (2016). Expansion Microscopy with Conventional Antibodies and Fluorescent Proteins. Nat. Methods 13, 485–488. 10.1038/nmeth.3833.

101. Asano, S.M., Gao, R., Wassie, A.T., Tillberg, P., Chen, F., and Boyden, E.S. (2018). Expansion Microscopy: Protocols for Imaging Proteins and RNA in Cells and Tissues. Curr. Protoc. Cell Biol. 80, e56. 10.1002/cpcb.56.

102. Gambarotto, D., Zwettler, F.U., Le Guennec, M., Schmidt-Cernohorska, M., Fortun, D., Borgers, S., Heine, J., Schloetel, J.-G., Reuss, M., Unser, M., et al. (2019). Imaging cellular ultrastructures using expansion microscopy (U-ExM). Nat. Methods 16, 71–74. 10.1038/s41592-018-0238-1.

103. Phan, S., Boassa, D., Nguyen, P., Wan, X., Lanman, J., Lawrence, A., and Ellisman, M.H. (2016). 3D reconstruction of biological structures: automated procedures for alignment and reconstruction of multiple tilt series in electron tomography. Adv. Struct. Chem. Imaging 2, 8. 10.1186/s40679-016-0021-2.

104. Ester, M., Kriegel, H.-P., Sander, J., and Xu, X. A Density-Based Algorithm for Discovering Clusters in Large Spatial Databases with Noise.

105. Heydarian, H., Schueder, F., Strauss, M.T., van Werkhoven, B., Fazel, M., Lidke, K.A., Jungmann, R., Stallinga, S., and Rieger, B. (2018). Template-free 2D particle fusion in localization microscopy. Nat. Methods 15, 781–784. 10.1038/s41592-018-0136-6.

106. Veatch, S.L., Machta, B.B., Shelby, S.A., Chiang, E.N., Holowka, D.A., and Baird, B.A. (2012). Correlation Functions Quantify Super-Resolution Images and Estimate Apparent Clustering Due to Over-Counting. PLOS ONE 7, e31457. 10.1371/journal.pone.0031457.

107. Schindelin, J., Arganda-Carreras, I., Frise, E., Kaynig, V., Longair, M., Pietzsch, T., Preibisch, S., Rueden, C., Saalfeld, S., Schmid, B., et al. (2012). Fiji: an opensource platform for biological-image analysis. Nat. Methods 9, 676–682. 10.1038/nmeth.2019.

108. Lord, S.J., Velle, K.B., Mullins, R.D., and Fritz-Laylin, L.K. (2020). SuperPlots: Communicating reproducibility and variability in cell biology. J. Cell Biol. 219, e202001064. 10.1083/jcb.202001064.

