## Supplemental Figures for "TorsinA is essential for the timing and localization of neuronal nuclear pore complex biogenesis"

### Figure S1

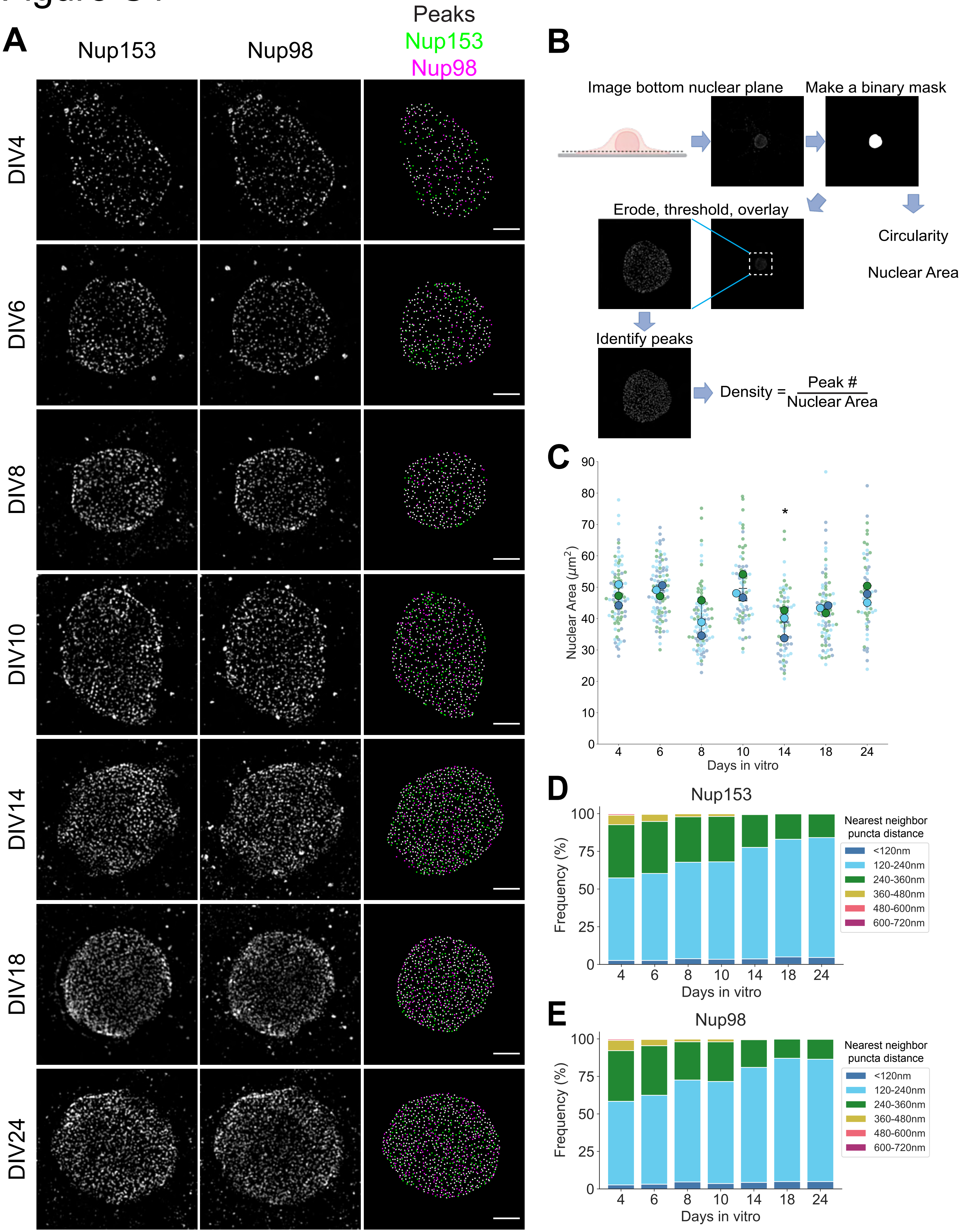

A
Nup153
Nup98
Peaks
Nup153
Nup98
DIV4
DIV6
DIV8
DIV10
DIV14
DIV18
DIV24
B
Image bottom nuclear plane
Make a binary mask
Erode, threshold, overlay
Circularity
Nuclear Area
Identify peaks
Density = Peak # / Nuclear Area
C
Nuclear Area (μm²)
Days in vitro
4 6 8 10 14 18 24
*
D
Nup153
Frequency (%)
Days in vitro
Nearest neighbor puncta distance
<120nm
120-240nm
240-360nm
360-480nm
480-600nm
600-720nm
E
Nup98
Frequency (%)
Days in vitro
Nearest neighbor puncta distance
<120nm
120-240nm
240-360nm
360-480nm
480-600nm
600-720nm

**Figure S1: Identification and analysis of nucleoporin puncta**

A) SIM images of primary neurons aged DIV4, 6, 8, 10, 14, 18 and 24 labeled with anti-Nup153 and anti-Nup98 antibodies. Peaks column shows peaks identified from Nup153 (green) and Nup98 (magenta) puncta. Scale bar = 2μm.

B) Schematic of image analysis pipeline. The flattest part of the nuclear envelope was imaged. Nuclear ROIs were determined by manually outlining each nucleus. ROIs were eroded to account for nuclear envelope curvature at the edge, then overlayed onto Otsu thresholded images. Local maxima were identified with extended-maxima transform and used to establish peaks corresponding to each Nup.

C) Superplots of nuclear ROI area as determined by Nup153/Nup98 staining in maturing neurons aged DIV4, 6, 8, 10, 14, 18 and 24. Plots show mean $\pm$ SD, with color coding indicative of biological replicates. Repeated measures one-way ANOVA with Dunnett's multiple comparisons test was performed, with DIV4 as the reference condition. Only DIV14 showed statistical significance; *P=0.0331.

D, E) Frequency distribution of nearest neighbor distances of Nup153 (D) and Nup98 (E) puncta. For each Nup153 and Nup98 peak, the distance to its nearest neighbor was calculated. The results from three biological replicates were combined for each timepoint.

### Figure S2

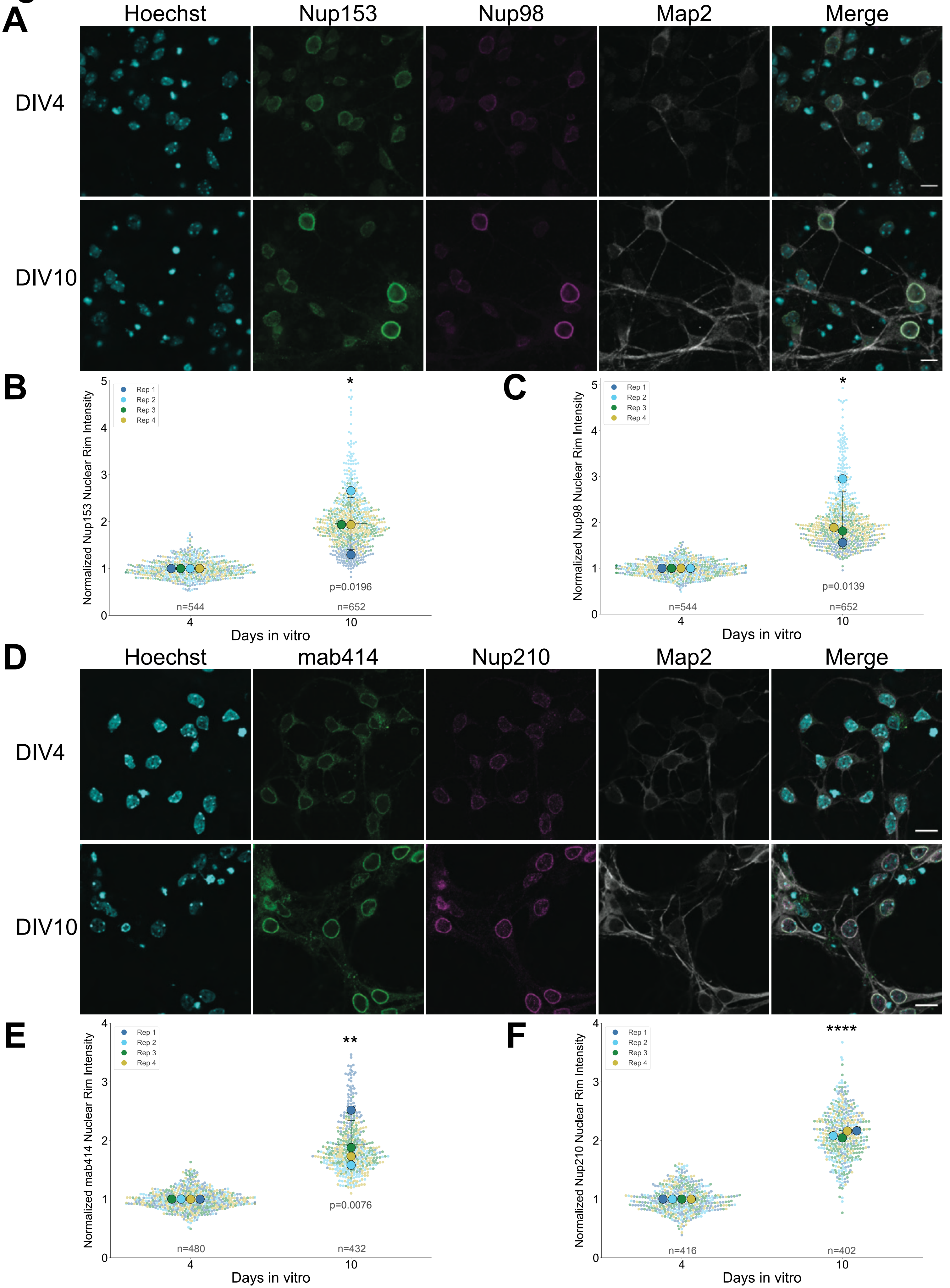
A
Hoechst
Nup153
Nup98
Map2
Merge
DIV4
DIV10
B
Rep 1
Rep 2
Rep 3
Rep 4
Normalized Nup153 Nuclear Rim Intensity
*
p=0.0196
n=544
n=652
4
10
Days in vitro
C
Normalized Nup98 Nuclear Rim Intensity
*
p=0.0139
n=544
n=652
Days in vitro
D
Hoechst
mab414
Nup210
Map2
Merge
DIV4
DIV10
E
Normalized mab414 Nuclear Rim Intensity
**
p=0.0076
n=480
n=432
Days in vitro
F
Normalized Nup210 Nuclear Rim Intensity
****
n=416
n=402
Days in vitro

**Figure S2: Multiple NPC components exhibit density increase during neuronal maturation**

A) Confocal images of DIV4 and DIV10 primary neurons labeled with anti-Nup153 and anti-Nup98 antibodies. Scale bar = 10μm.

B, C) Normalized nuclear rim fluorescence intensity of Nup153 and Nup98 in DIV4 and DIV10 primary neurons. Plots show mean $\pm$ SD, with color coding indicative of biological replicates. P values were calculated using paired t-test.

D) Confocal images of DIV4 and DIV10 primary neurons labeled with anti-FG nucleoporin (mab414) and anti-Nup210 antibodies. Scale bar = 10μm.

E, F) Normalized nuclear rim fluorescence intensity of FG-Nups (mab414) and Nup210 in DIV4 and DIV10 primary neurons. Plots show mean $\pm$ SD, with color coding indicative of biological replicates ****$P<0.0001$, paired t-test.

### Figure S3

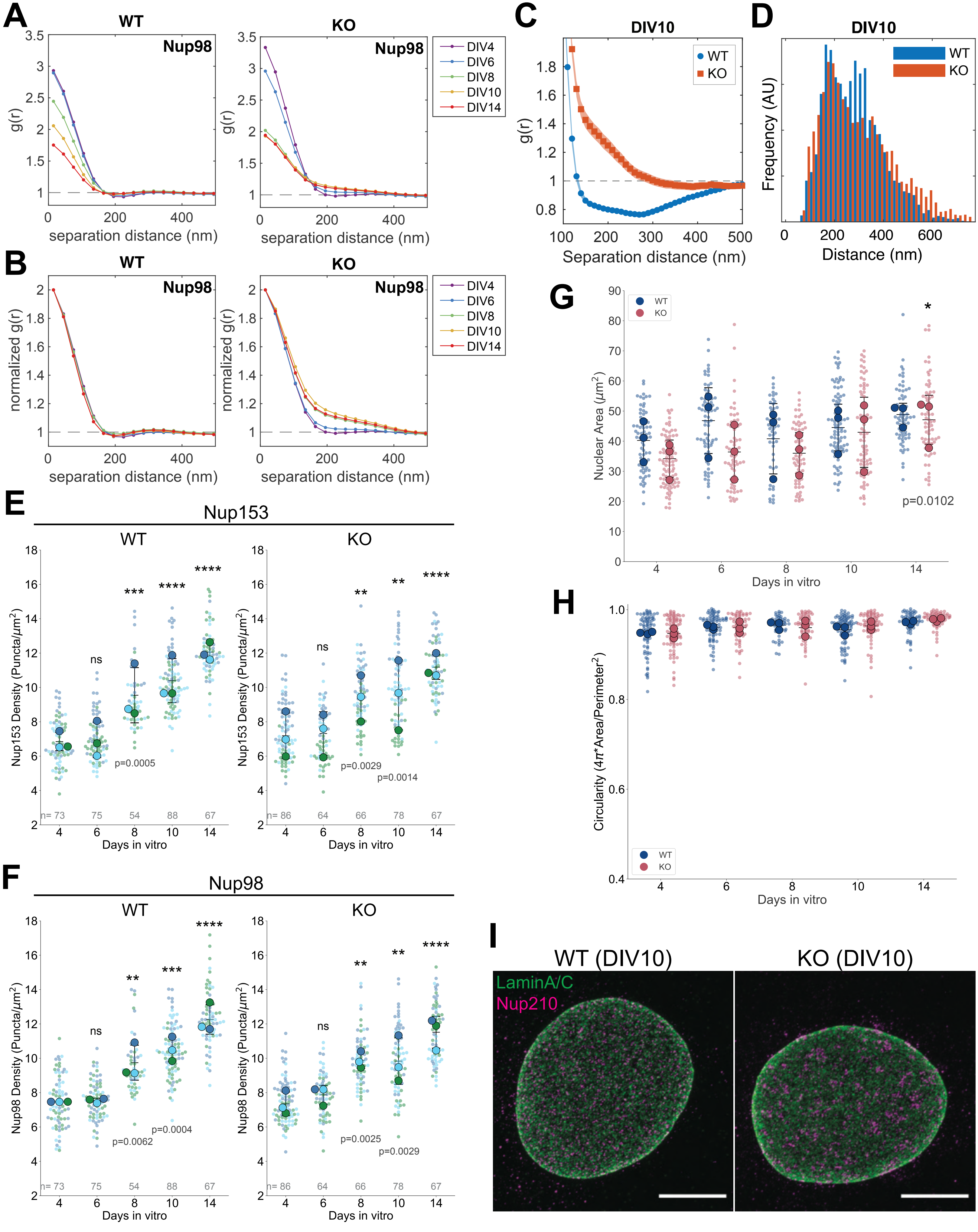

A
WT
KO
Nup98
g(r)
separation distance (nm)
DIV4
DIV6
DIV8
DIV10
DIV14
B
normalized g(r)
C
DIV10
WT
KO
Separation distance (nm)
D
Frequency (AU)
Distance (nm)
E
Nup153
Nup153 Density (Puncta/μm²)
Days in vitro
ns
***
****
**
p=0.0005
p=0.0029
p=0.0014
n= 73
75
54
88
67
n= 86
64
66
78
F
Nup98 Density (Puncta/μm²)
p=0.0062
p=0.0004
p=0.0025
p=0.0029
G
Nuclear Area (μm²)
p=0.0102
H
Circularity (4π*Area/Perimeter²)
I
WT (DIV10)
KO (DIV10)
LaminA/C
Nup210

**Figure S3: Comparison of WT and torsinA-KO neuronal NPC spatial organization and nuclear morphology**

A) Autocorrelation plot of WT and torsinA-KO neurons calculated from Nup98 WT and torsinA-KO SIM images.

B) Normalized autocorrelation plot of WT and torsinA-KO neurons calculated from Nup98 WT and torsinA-KO SIM images. Autocorrelations were normalized to account for amplitude dependency on NPC density.

C) Autocorrelation plot of DIV10 WT and torsinA-KO neurons calculated from Nup210 dSTORM images.

D) Nearest neighbor distance of segmented DIV10 WT and torsinA-KO NPC centroids.

E, F) WT and torsinA-KO Nup153 (E) and Nup98 (F) puncta density in maturing primary neurons aged DIV4, 6, 8, 10, 14. Plots show mean $\pm$ SD, with color coding indicative of biological replicates. Timepoints from the same biological replicate were matched. ns, non-significant; ****$P<0.0001$; repeated measures two-way ANOVA with Dunnett's multiple comparisons test, using DIV4 as a reference.

G) Nuclear area of WT and torsinA-KO neurons calculated from manually drawn ROIs from SIM images. Plots show mean $\pm$ SD. Repeated measures two-way ANOVA with Sidak's multiple comparisons test was used to compare genotypes at each timepoint, and no comparisons reached statistical significance. Repeated measures two-way ANOVA with Dunnett's multiple comparisons test was used to compare all timepoints to DIV4 within each genotype.

H) Nuclear ROI circularity of WT and torsinA-KO neurons calculated from manually drawn ROIs from SIM images. Plots show mean $\pm$ SD. Repeated measures two-way ANOVA with Sidak's multiple comparisons test was used to compare genotypes at each timepoint. No statistical significance was detected.

I) Expansion microscopy of DIV10 WT and torsinA-KO neurons labeled with Nup210 and Lamin A/C antibodies. Scale bar = 20µm.

### Figure S4

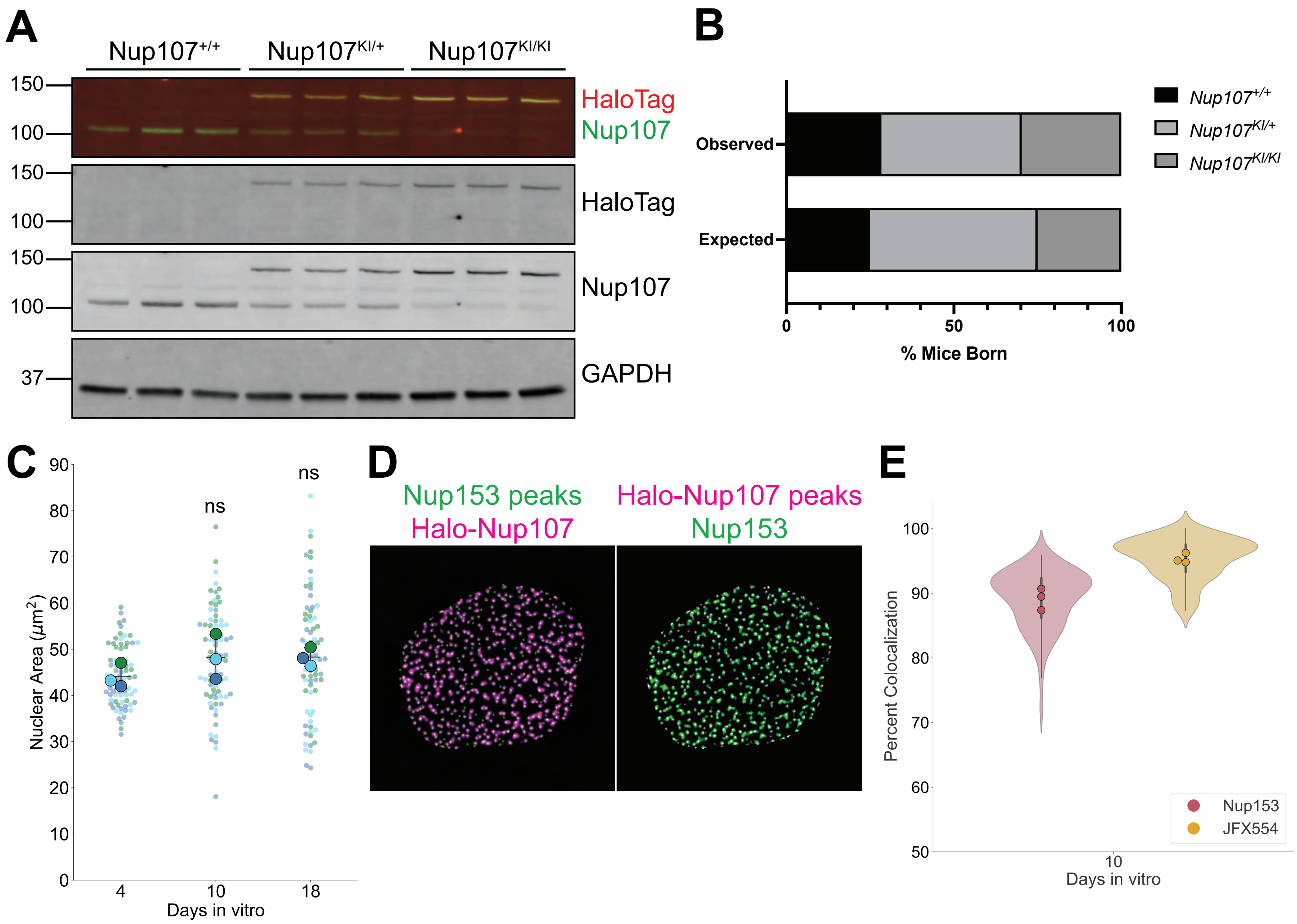

A
Nup107+/+
Nup107KI/+
Nup107KI/KI
150
100
HaloTag
Nup107
150
100
HaloTag
150
100
Nup107
37
GAPDH
B
Observed
Expected
0
50
100
% Mice Born
Nup107+/+
Nup107KI/+
Nup107KI/KI
C
Nuclear Area (μm2)
0
10
20
30
40
50
60
70
80
90
ns
ns
4
10
18
Days in vitro
D
Nup153 peaks
Halo-Nup107
Halo-Nup107 peaks
Nup153
E
Percent Colocalization
50
60
70
80
90
100
Nup153
JFX554
10
Days in vitro

**Figure S4: Validation of HaloTag-Nup107 mouse line**

A) Immunoblot of P0 cortical lysates from *Nup107*$^{+/+}$, *Nup107*$^{KI/+}$, and *Nup107*$^{KI/KI}$ mice probed with anti-Nup107 and anti-HaloTag antibodies. GAPDH was blotted as a loading control. Each band represents an independent biological sample (3 animals for each genotype).

B) Distribution of genotypes in litters derived from intercrossing *Nup107*$^{KI/+}$ mice. No deviation from the expected Mendelian ratio was observed (P=0.3729, Chi-squared test). 74 pups from 9 litters were analyzed.

C) Superplots of nuclear ROI area from Nup153 and JFX554 SIM images. Plots show mean ± SD, with color coding indicative of biological replicates. ns, not significant; repeated measures one-way ANOVA with Dunnett's multiple comparisons test, with DIV4 as a reference condition.

D) Representative image of Nup153 and HaloTag-Nup107 colocalization in DIV10 HaloTag-Nup107 neurons. Identified Nup153 peaks are overlaid with thresholded JFX554 SIM image and vice versa.

E) Violin plot showing the percent of Nup153 and JFX554 puncta that colocalize with JFX554 and Nup153, respectively, in DIV10 neurons. Nup153 colocalization with JFX554 was measured by calculating the % of Nup153 peaks that overlap with thresholded JFX554 puncta. Conversely, JFX554 colocalization with Nup153 was measured by calculating the % of JFX554 peaks that overlap with thresholded Nup153 puncta.

### Figure S5

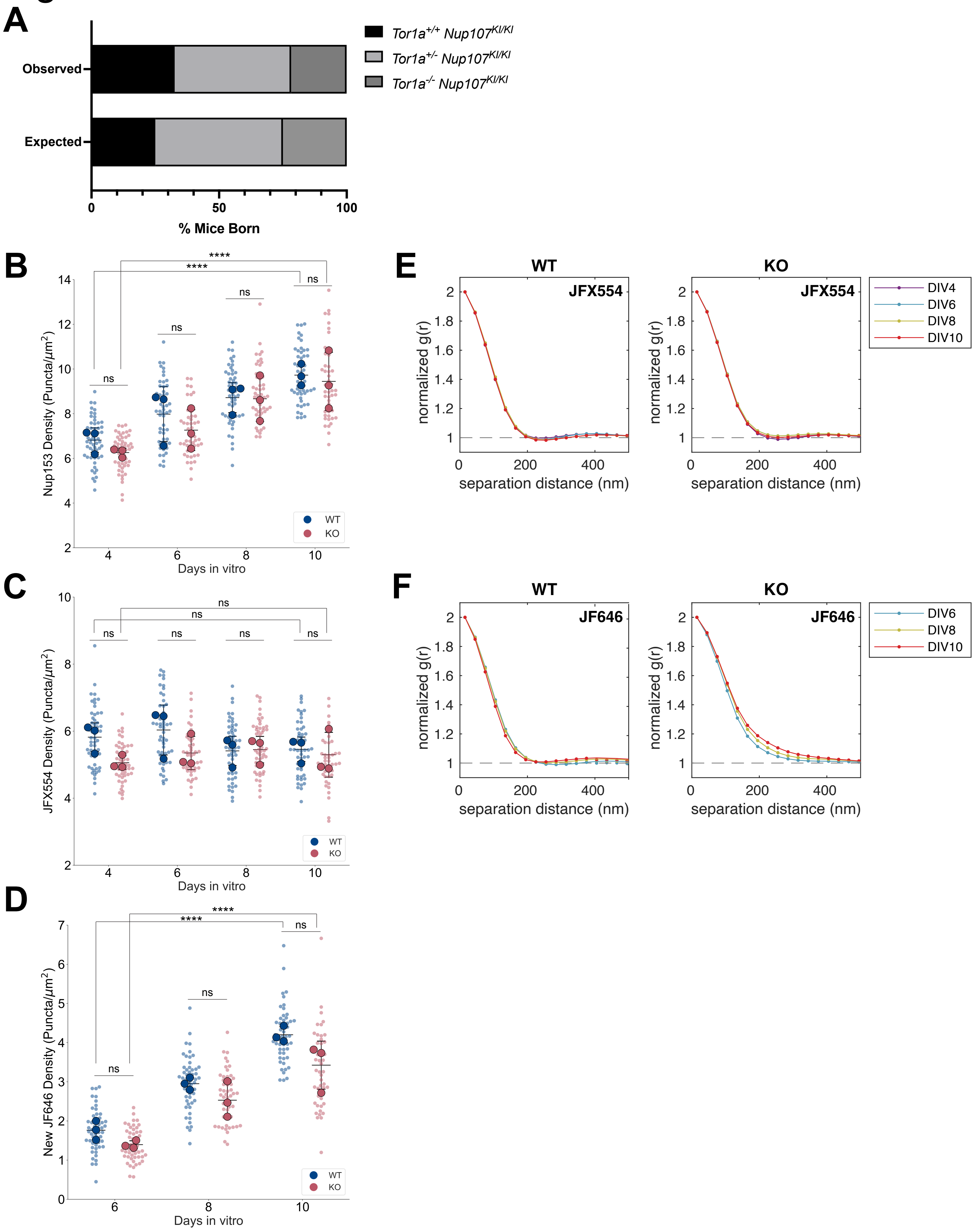
A
Observed
Expected
0
50
100
% Mice Born
Tor1a+/+ Nup107KI/KI
Tor1a+/- Nup107KI/KI
Tor1a-/- Nup107KI/KI
B
Nup153 Density (Puncta/µm2)
Days in vitro
ns
****
WT
KO
C
JFX554 Density (Puncta/µm2)
Days in vitro
D
New JF646 Density (Puncta/µm2)
Days in vitro
E
WT
KO
JFX554
normalized g(r)
separation distance (nm)
DIV4
DIV6
DIV8
DIV10
F
JF646

**Figure S5: Comparison of NPC density and distribution from HaloTag-Nup107 pulse-chase**

A) Distribution of genotypes in litters derived from intercrossing *Tor1a*$^{+/-}$*; Nup107*$^{KI/KI}$ mice. All genotypes were born at the expected Mendelian ratio (P=0.4880, Chi-squared test). 46 pups from 6 litters were analyzed.

B, C, D) Nup153 (B), JFX554 (C), and new JF646 (D) puncta density in maturing WT and torsinA-KO primary neurons aged DIV4, 6, 8, 10. Plots show mean $\pm$ SD. Repeated measures two-way ANOVA with Sidak's multiple comparisons test was used to compare genotypes at each timepoint, and no comparisons reached statistical significance. Repeated measures two-way ANOVA with Dunnett's multiple comparisons test was used to compare all timepoints to DIV4, with ****P<0.0001.

E) Normalized autocorrelation of JFX554 SIM images over 0-500nm separation distance. Autocorrelation values <1 around 200nm in both WT and torsinA-KO neurons reflect nonrandom uniform distribution of JFX554-labeled NPCs.

F) Normalized autocorrelation of JF646 SIM images over 0-500nm separation distance. Autocorrelation values >1 in torsinA-KO neurons reflect clustering of NPCs labeled with JF646.

### Figure S6

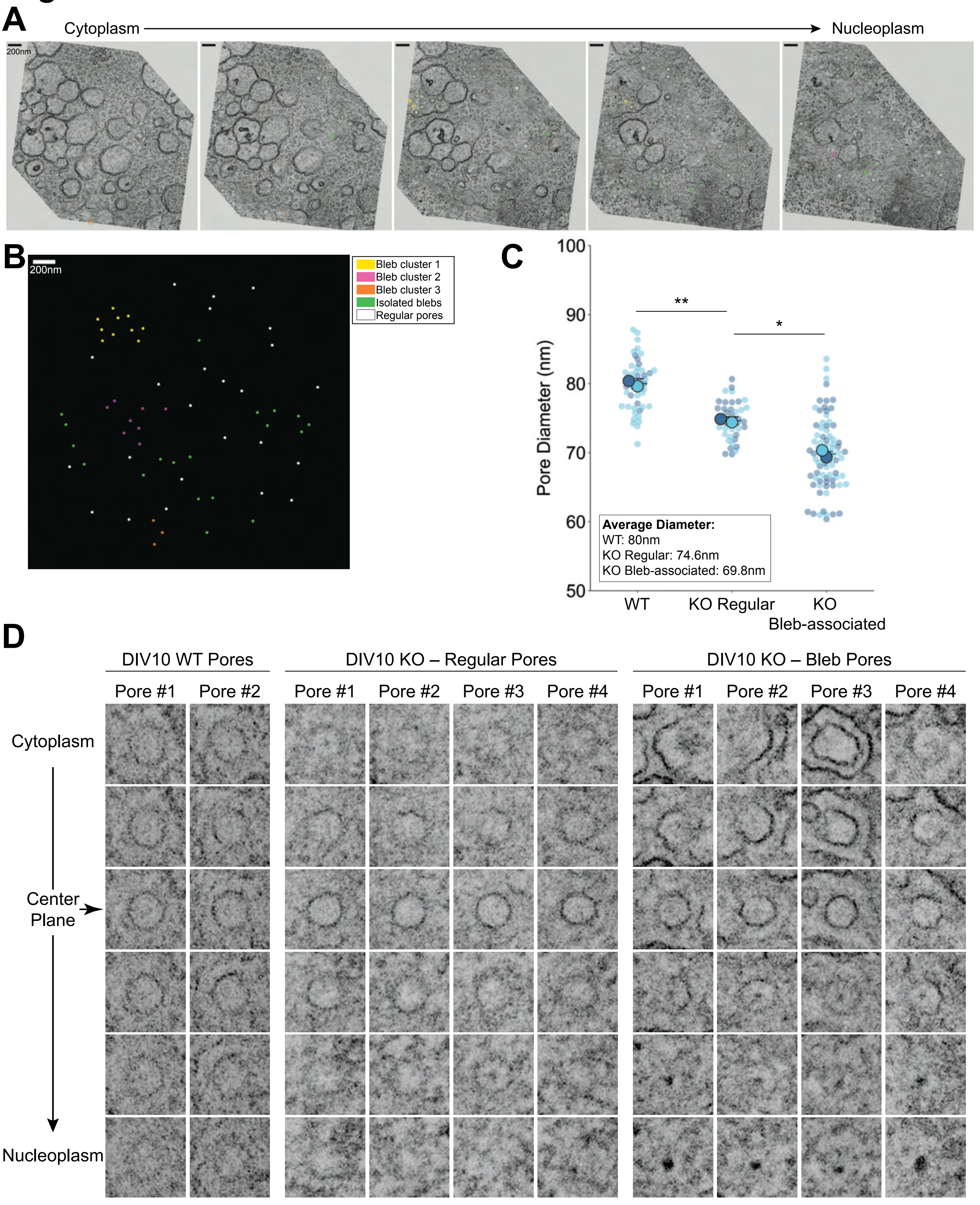
A
Cytoplasm
Nucleoplasm
200nm
B
200nm
Bleb cluster 1
Bleb cluster 2
Bleb cluster 3
Isolated blebs
Regular pores
C
Pore Diameter (nm)
100
90
80
70
60
50
**
*
Average Diameter:
WT: 80nm
KO Regular: 74.6nm
KO Bleb-associated: 69.8nm
WT
KO Regular
KO Bleb-associated
D
DIV10 WT Pores
DIV10 KO – Regular Pores
DIV10 KO – Bleb Pores
Pore #1
Pore #2
Pore #3
Pore #4
Cytoplasm
Center Plane
Nucleoplasm

**Figure S6: Bleb-associated pores are narrow and exhibit nucleoplasmic central plugs**

A) Slices from a DIV10 torsinA-KO tomogram oriented as an en-face view of the neuronal nuclear membrane. NPCs localized to clusters of blebs, isolated blebs, or regular NPCs without blebs are marked by color-coded spheres centered around the middle plane of the pore channel, denoted by larger circles.

B) Projection of all marked pores from the tomogram shown in (A).

C) Superplot of WT, KO regular (non bleb-associated), and KO bleb-associated pore diameter (nm). Plots show mean ± SD, with color coding indicative of biological replicates. Swarmplot of individual pore diameters (small points) is overlayed with the mean of pore diameters for each biological replicate (larger points). Pores from two WT and two torsinA-KO tomograms were analyzed. KO regular and bleb-associated pores were identified from the same torsinA-KO tomograms. *P=0.0128, **P=0.0068; unpaired t-test.

D) Slices showing individual pores from DIV10 WT and torsinA-KO tomograms. Regular and bleb-associated torsinA-KO pores were sampled from the same tomogram. From top to bottom, the slices progress from the cytoplasmic side of the pore channel towards the nucleoplasm. Center plane of the pore channel is marked with a horizontal arrow.

### Figure S7

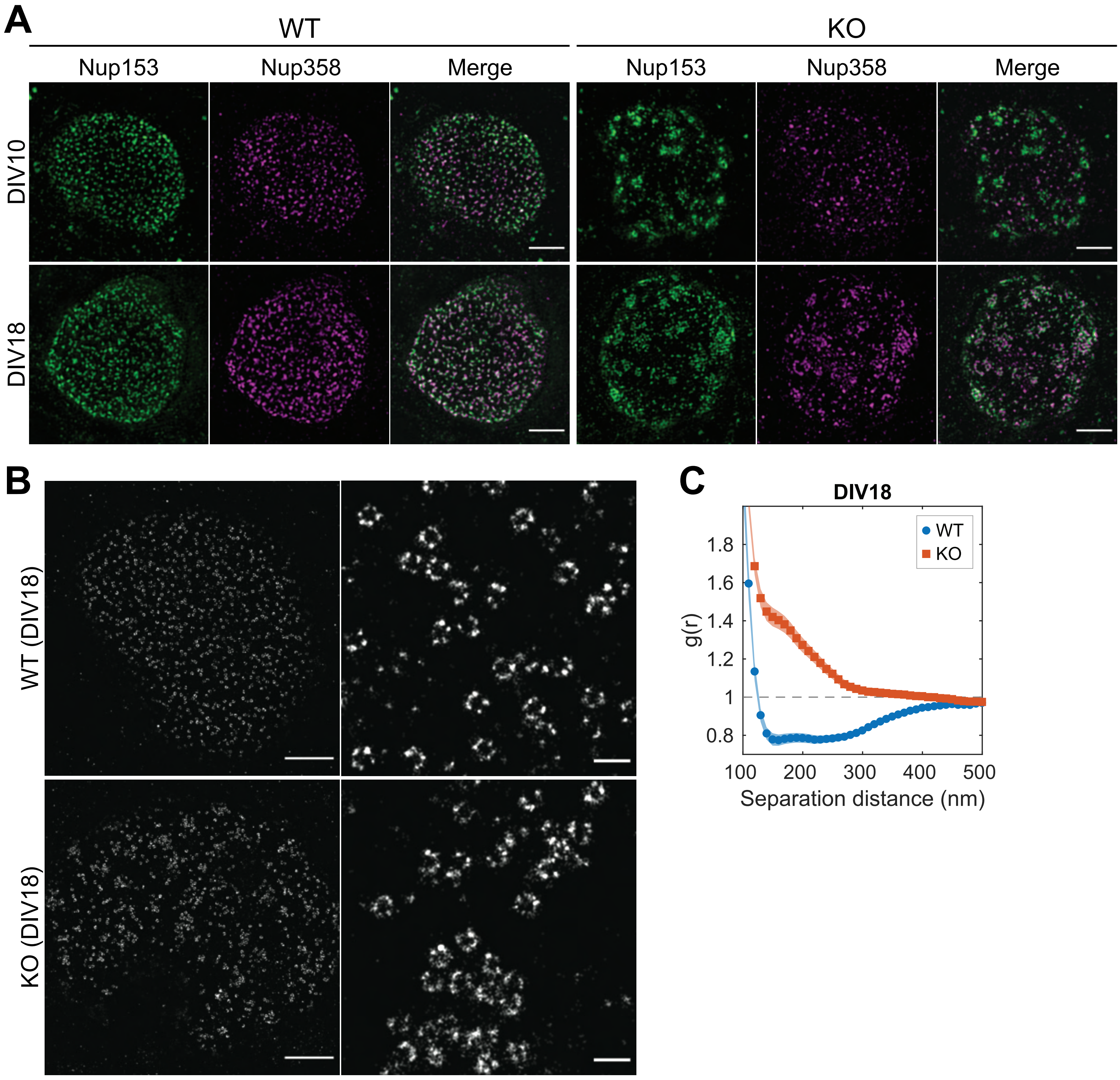
A
WT
KO
Nup153
Nup358
Merge
DIV10
DIV18
B
WT (DIV18)
KO (DIV18)
C
DIV18
WT
KO
g(r)
1.8
1.6
1.4
1.2
1
0.8
100
200
300
400
500
Separation distance (nm)

**Figure S7: Nup358 is recruited to persisting NPC clusters in DIV18 torsinA-KO neurons**

A) SIM images of DIV10 and DIV18 WT and torsinA-KO neurons labeled with Nup153 and Nup358 antibodies. Scale bar = 2μm.

B) dSTORM images of Nup210 in DIV18 WT and torsinA-KO neurons. Scale bar = 2nm. Right panels show zoomed in view. Scale bar for right panels = 200nm.

C) Autocorrelation plot of DIV18 WT and torsinA-KO neurons, determined from Nup210 dSTORM images. 18 WT and 17 KO cells from 3 biological replicates were analyzed.
